# The spatial transcriptomic landscape of the healing intestine following damage

**DOI:** 10.1101/2021.07.01.450768

**Authors:** Sara M. Parigi, Ludvig Larsson, Srustidhar Das, Ricardo O. Ramirez Flores, Annika Frede, Kumar P. Tripathi, Oscar E. Diaz, Katja Selin, Rodrigo A. Morales, Xinxin Luo, Gustavo Monasterio, Camilla Engblom, Nicola Gagliani, Julio Saez-Rodriguez, Joakim Lundeberg, Eduardo J. Villablanca

## Abstract

The intestinal barrier is composed of a complex cell network defining highly compartmentalized and specialized structures. Here, we use spatial transcriptomics (ST) to define how the transcriptomic landscape is spatially organized in the steady state and healing murine colon. At steady state conditions, we demonstrate a previously unappreciated molecular regionalization of the colon, which dramatically changes during mucosal healing. Here, we identified spatially-organized transcriptional programs defining compartmentalized mucosal healing, and regions with dominant wired pathways. Furthermore, we showed that decreased p53 activation defined areas with increased presence of proliferating epithelial stem cells. Finally, we used our resource to map transcriptomics modules associated with human diseases demonstrating that ST can be used to inform clinical practice. Overall, we provide a publicly available resource defining principles of transcriptomic regionalization of the colon during mucosal healing and a framework to develop and progress further hypotheses.

## Introduction

The intestine is divided into the small and large bowels that together host the highest density of commensal microbiota, which in turn is spatially heterogeneous across the proximal-distal axis ^1^. The geographically heterogeneous microbial exposure has contributed to the establishment of a highly compartmentalized organ that has distinct functions depending on the proximal-distal location ^2,3^. For example, vitamin A-metabolizing enzymes and consequently retinoic acid production and function are higher in the proximal compared to distal small intestine ^4^, generating a proximal-to-distal gradient. Although it is broadly accepted that the small intestine is highly compartmentalized, whether a clear molecular regionalization exists in the colon is yet to be determined.

The intestine relies on the constant regeneration of the intestinal epithelium to maintain homeostasis. Breakdown in regenerative pathways may lead to pathogen translocation and the development of chronic intestinal pathologies, such as inflammatory bowel disease (IBD) ^5^. Therefore, the intestinal barrier must quickly adapt to promote tissue regeneration and healing following injury. However, the cellular and molecular circuitry at steady state conditions and how it adapts upon challenge is yet to be fully characterized.

The intestine offers a unique opportunity to investigate common principles of tissue repair at the barrier because of its spatial organization, which is fundamental to its function. When intestinal barrier injury occurs, damaged epithelial cells are shed and replaced by mobilizing intestinal stem cell (ISC)-derived cells ^6^, a phenomenon highly dependent on signals coming from the neighboring microenvironment (niche) ^7^. Similarly, immune cells are recruited or expanded *in situ* to protect the host from invading pathogens and to orchestrate the healing process by providing resolving signals ^8^. Although initially considered as a mere structural support, stromal cells, which includes fibroblasts, endothelial/lymphatic cells, pericytes and glial cells, are also actively involved in barrier healing through tissue remodeling, matrix deposition, neoangiogenesis, muscle contraction, and production of pro-regenerative signals ^9^. Therefore, immune, epithelial and stromal cells must quickly adapt within a defined microenvironment and establish a molecular network to promote tissue repair. However, whether different segments of the intestine and their microenvironments possess distinct types of tissue repair mechanisms is currently unknown.

Our previous study unveiled the temporal transcriptomic dynamic of the colonic tissue over the course of dextran sodium sulphate (DSS) colitis, identifying genes and pathways differentially modulated during acute injury or regeneration ^10^. Although bulk or single cell RNA sequencing studies provide unbiased transcriptome analysis, the spatial context within the tissue is typically lost. In contrast, targeted technologies for spatial gene expression analysis (e.g. in situ RNA-sequencing, fluorescence in situ hybridization [FISH], RNA-scope) require knowledge of specific candidate genes to interrogate and thus do not allow an unsupervised investigation of pathways enriched in healing areas.

To overcome these limitations, we exploited spatial transcriptomics (ST), an unbiased technology allowing sequencing of polyadenylated transcripts from a tissue section, which can be spatially mapped onto the histological brightfield image ^11^. ST allowed us to uncover an unprecedented view of the molecular regionalization of the murine colon, which was further validated on human intestinal specimens. By comparing ST of colonic tissue under steady state and upon mucosal healing (i.e. from DSS-treated mice), we identified and spatially mapped transcriptional signatures of tissue repair processes, immune cell activation/recruitment, pro-regenerative pathways, and tissue remodeling. The spatial landscape of pathway activity during mucosal healing also unveiled a negative correlation between p53 activity and proliferating epithelial stem cells. Moreover, targeted mapping of genes associated with disease outcome in human IBD patients and IBD risk variants identified from genome-wide association studies (GWAS) allowed us to infer their involvement in specific pathological processes based on their localization in areas with distinct histological properties.

## Results

### Spatial transcriptomics revealed distinct molecular regionalization of murine colonic epithelium at steady state condition

To characterize the transcriptomic landscape of the colon tissue at steady state condition, we processed frozen colons for ST using the Visium (10X Genomics) platform (Fig. 1A). The pre-filtered dataset corresponded mostly to protein coding genes (Extended Data Fig. 1A). Upon filtering out non-coding RNAs (ncRNAs) and mitochondrial protein coding genes, the resulting dataset consists of 2604 individual spots, with an average of ∼4125 genes and ∼11801 unique transcripts per spot (Extended Data Fig. 1B). First, we deconvolved the spatial transcriptomic dataset using non-negative matrix factorization (NNMF) to infer activity maps ^12^, and we restricted the analysis to only 3 factors that capture the most basic structure of the colon at steady state conditions (d0) (Fig. 1B). We identified 3 basic structural transcriptomic landscapes that were histologically discernible as intestinal epithelial cells (IEC) (NNMF_3), muscle (NNMF_2), and a mixture between lamina propria (LP) and IEC (NNMF_1), which were indistinguishable from the IEC towards the most distal colon (Fig. 1B). Analysis of the top contributing genes for each factor confirmed the identity of the muscle and IEC, and the mixed signature between IEC, muscle, and LP (Fig. 1C). Using immunohistochemistry (IHC) data from the human protein atlas ^13^, we validated the specific expression of CDH17 (also known as liver-intestine cadherin or LI cadherin)^14^ and TAGLN (transgelin, smooth muscle marker) in the IEC and muscularis layer, respectively (Fig. 1D). By contrast, ADH1 (alcohol dehydrogenase 1) showed a mixed expression between the LP and IEC compartments (Fig. 1D). To better visualize the molecular regionalization both across the proximal-distal and serosa-luminal axis, we digitally unrolled the colon (Extended Data Fig. 2A and Fig. 1E) as described in methods. In line with the ST expression in Fig. 1C, the muscle, LP/IEC and proximal IEC factors were enriched in the corresponding regions of the distal-proximal and serosa-luminal axis of the digitally unrolled colon (dark dots in Fig. 1E). Among the genes driving factor 3 (NNMF_3) we found *Car1, Mettl7b, Emp1, Fabp2*, and *Hmgcs2* that were highly expressed in the proximal colon (Fig. 1F and Extended Data Fig. 2B). Our data aligns with reports showing *Car1* promoter-driven expression in the proximal but not distal colon ^15^. In contrast, *Retnlb, Sprr2a2* and *Ang4* were enriched in the mid colon, and *Prdx6, Tgm3, Ly6g, Eno3, and B4galt1* were enriched in the distal colon (Fig. 1F and Extended Data Fig. 2B). Because they have not been previously described as markers for the distinct colonic compartments, we used qPCR to validate the region-specific mRNA expression of genes coding for the ketogenic rate-limiting enzyme, mitochondrial 3-hydroxy-3-methylglutaryl-CoAsynthase 2 (HMGCS2), the antimicrobial peptide angiogenin 4 (ANG4), and the Beta-1,4-galactosyltransferase 1 (B4GALT1)(Fig. 1G). Therefore, ST analysis distinguished a stratification between the LP, IEC, and muscle compartment which is clearly evident at the proximal but not distal colon.

**Fig. 1.**
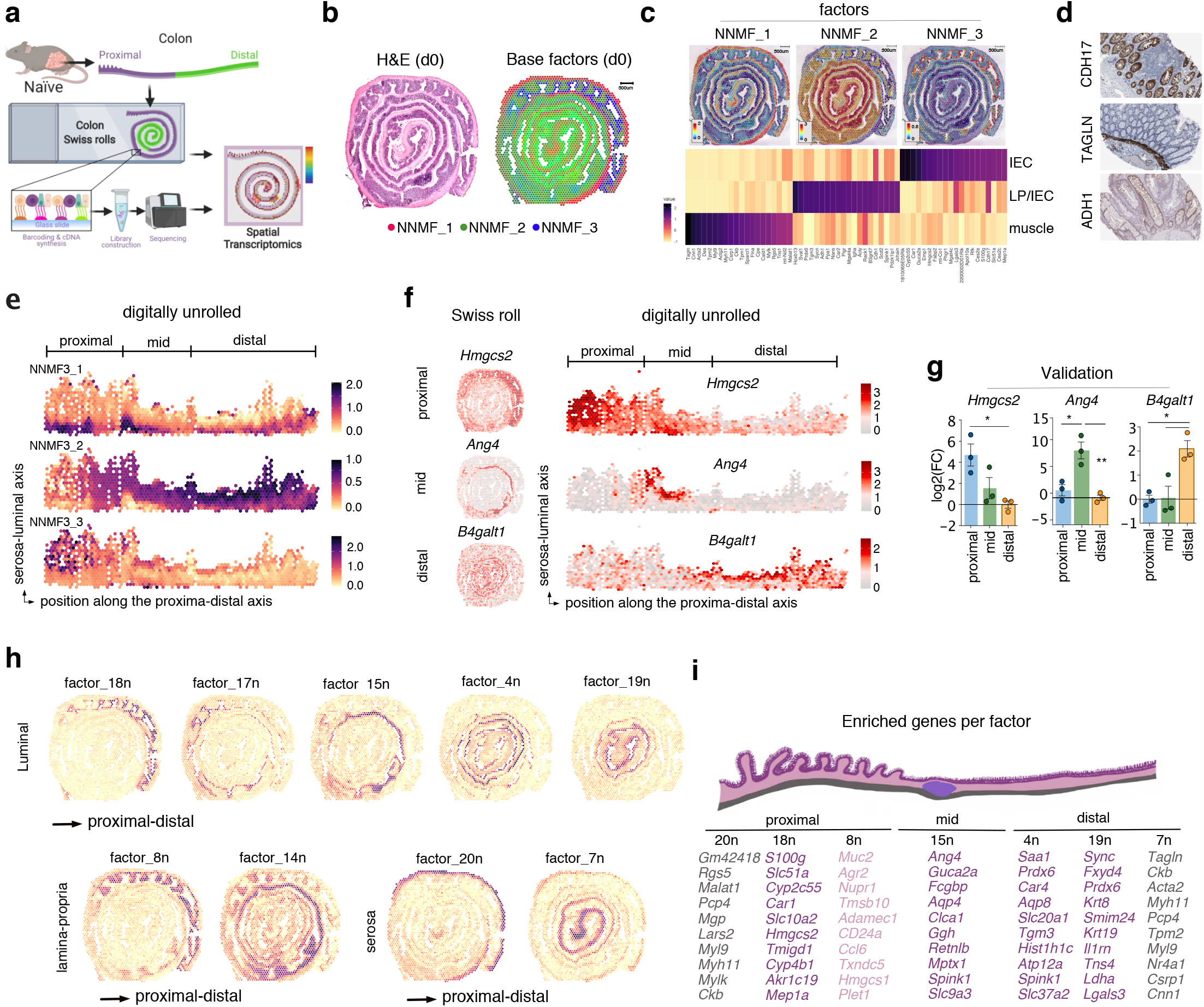
Spatial transcriptomics reveals molecular regionalization of the murine colonic tissue in steady state. (A) Schematics of the experimental design: the colonic tissue from a naive wild type mouse was processed as a Swiss roll for spatial transcriptomic (ST) with VISIUM 10X technology. (B) Colon Swiss rolls shown in hematoxylin and eosin staining (left) and with each ST spot color coded based on non-negative matrix factorization (NNMF) (right). ST spots belonging uniquely to one factor are colored in red, blue and green for NNMF1, 2 and 3 respectively. ST spots shared between different factors are colored with respective intermediate gradation of these 3 colors. (C) Top: spatial distribution of the 3 factors distinguishing muscle, lamina propria (LP) and intestinal epithelial cells (IEC). Bottom: heatmap showing the top 20 genes defining each factor. (D) Immunohistochemical staining of CDH17, TAGLN, ADH1 in healthy human colonic tissue (from Human Protein Atlas). (E) Digitally unrolled colonic tissue, showing the distribution of the 3 factors from Fig. 1C along the serosa-luminal and proximal-distal axis. (F) Proximal to distal distribution of *Hmgcs2, Ang4* and *B4galt1* expression in colonic swiss rolls (left) and digitally unrolled colon (right). (G) qPCR validation of regional expression of *Hmgcs2, Ang4* and *B4galt1* in proximal, mid and distal colonic biopsies from wild type mice (n=3, each dot represents one mouse). (H) Spatial distribution of 9 out of 20 factors in the naive colon displaying transcriptional regionalization along the serosa-luminal and proximal-distal axis. Each ST spot is assigned a color-coded score based on the expression of the genes defining each factor. (I) Schematic representation of the colon (top) and top genes annotated in Factors describing molecular regionalization of the naive colon (bottom). Factors are grouped based on their proximal-distal distribution and color-coded (i.e. grey-pink-purple) based on their serosa-luminal distribution.

Some studies have shown structural and functional differences between proximal and distal colon, specifically with respect to the epithelium ^15-20^; however, systematic and unsupervised molecular regionalization of the colon is lacking. To objectively identify genes differentially expressed in specific compartments of the colon, we used the NNMF method to distinguish relevant sources of variability of the data ^12^. Detailed factor analyses of the naive colon (denoted with “n”) resulted in a more pronounced/apparent colonic compartmentalization, in which the top genes defining a factor were sufficient to specifically demarcate the regionalization in the colon (Extended Data Fig. 3A-B). In particular, these factors defined a proximal-distal and a serosa-luminal axis (Fig. 1H). Functional enrichment analysis using the top contributing genes of factors defining the proximal and distal colonic IEC suggested that the murine proximal colon is specialized in water absorption, while the distal colon is specialized in solute transport (Extended Data Fig. 3C), indicating functional differences between the proximal and distal IEC compartments. Altogether, ST permitted the identification of a previously unappreciated level of colonic molecular compartmentalization in the steady state colon, as summarized in Fig. 1I.

### Visualization of lymphoid structures by factors enriched with B-cell associated genes

Next, we examined the capacity of our dataset to resolve macroscopic structures, such as lymphoid clusters, within the tissue. We observed an enrichment of B cell-associated genes in factors 1n, 3n and 9n (Fig. 2A). Upon mapping these factors onto the colonic tissue, we observed that factors 1n and 3n defined structures that resembled lymphoid aggregates, known as isolated lymphoid follicles (ILF) and/or cryptopatches (CP) (Fig. 2B). In contrast, factor 9n defined the colonic LP and was characterized by high expression of genes such as *Igha, Jchain, Igkc* characteristic of plasma cells (Fig. 2B). Among top-listed genes found in factor_3n, we validated *Clu* protein expression in lymphoid follicles ^21^ (Fig. 2C). Similarly, expression of JCHAIN, a small 15 kDa glycoprotein produced by plasma cells that regulates multimerization of secretory IgA and IgM and facilitates their transport across the mucosal epithelium ^22^, was validated by IHC in the human colonic LP (Fig. 2C). Pathways analysis confirmed that these factors were associated with immune responses (Fig. 2D). Whereas pathways associated with factor 3n suggested sites of lymphocyte priming (defined by processes associated with lymphocyte activation), pathways associated with factor 9n suggested sites of effector immune responses (defined by processes associated with adaptive immunity) (Fig. 2D). Interestingly, factor_1n, which defined the cells/region overlying the ILF, was characterized by genes such as *Ccl20* known to recruit CCR6^+^ B cells ^23,24^ and *Il22ra2*/*Il22bp*, a soluble receptor that neutralizes the effects of IL-22, a pleiotropic cytokine primarily expressed by lymphoid tissue inducer cells (LTis) ^25^. In addition, factor_1n was associated with pathways involved in cell mobilization and response to external stimulus (Fig. 2D). Because of the observation that factor 1n defines ILFs, it is tempting to propose that factor 1n-defined structures may serve as an anlagen for further maturation into ILF (factor_3n).

**Fig. 2.**
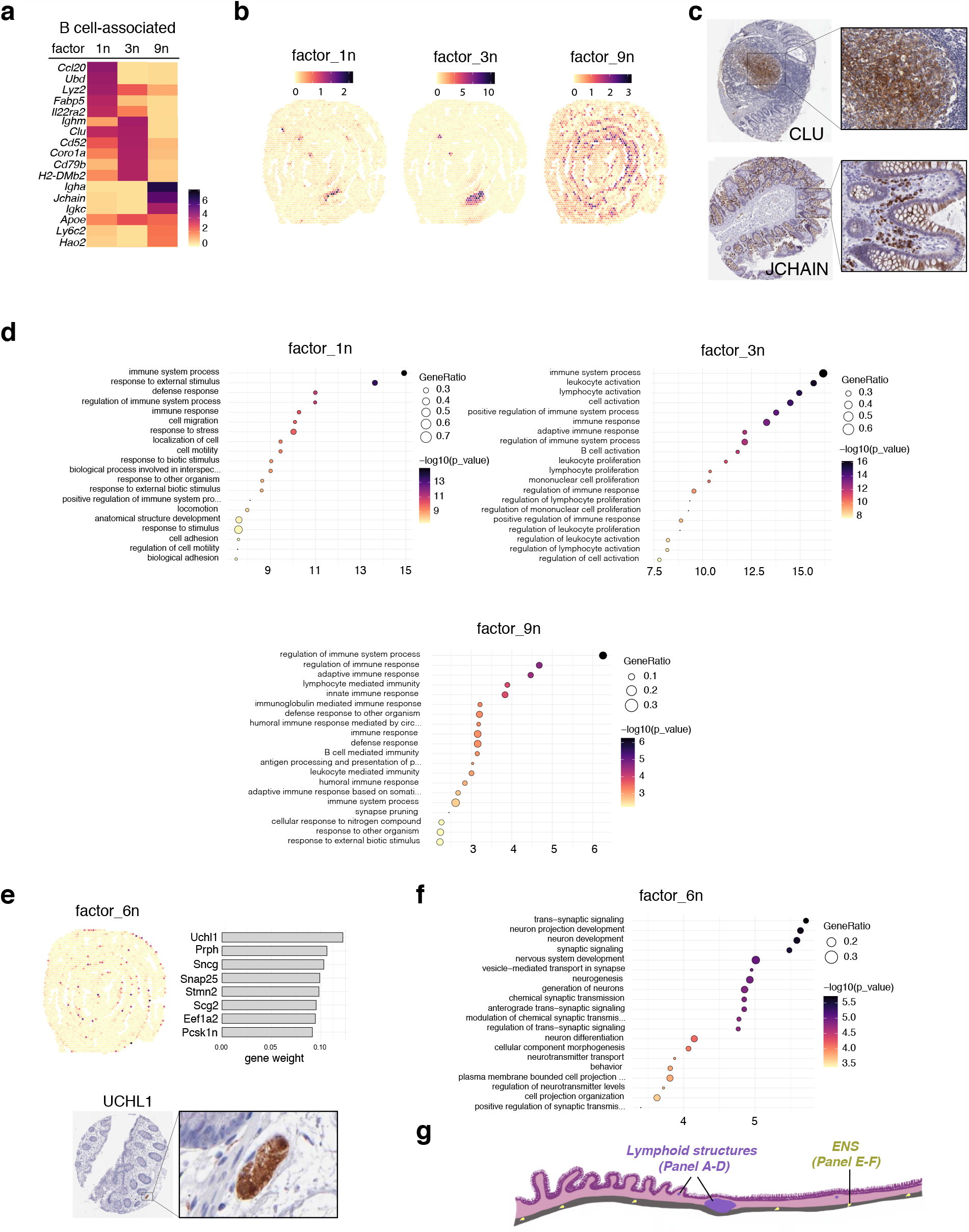
Identification and regional distribution of lymphoid follicle, B cell-associated, and enteric nervous system signatures in the naive murine colon. (A) Heatmap of the top genes defining factors 1n, 3n and 9n enriched in B cell signature. (B) Spatial distribution of B cell-associated factors in the naive colon. (C) Immunohistochemical staining of CLU (enriched in lymphoid follicles) and JCHAIN (localized in the lamina propria) in healthy human colonic tissue (from Human Protein Atlas). (D) Functional enrichment analysis (GO) of factors 1n, 3n and 9n. (E) Top: spatial distribution (left) and top genes (right) defining factor 6n (enteric nervous system). Bottom: immunohistochemical staining of UCHL1 (neuronal marker) in the colonic submucosa of healthy human colonic tissue (from Human Protein Atlas). (F) Pathway analysis (GO) of factor 6n. (G) Schematic representation of spatial distribution of B cell factors (i.e. 1n, 3n and 9n from Panels A-D) and ENS-factor 6n (from Panel E-F).

### Factor analysis identified molecular signatures that define areas associated with the enteric nervous system

Factor 6n was characterized by an enrichment of enteric nervous system (ENS)-associated genes which were located in the muscle area (Fig. 2E). Among the top-listed genes, we validated ubiquitin C-terminal hydrolase L1 (UCHL1) (Fig. 2E), which is specifically expressed in neurons ^26^. Functional enrichment analysis confirmed that factor_6n defined a transcriptomic profile associated with the ENS (Fig. 2F). In summary, the resolution of our Visium dataset permitted the identification of known structures (e.g. ENS and ILFs; Fig. 2G), thereby providing a platform to further investigate specific molecular circuitry within such regions.

### Molecular landscape of intestinal mucosal healing

Next, we sought to spatially resolve the colonic transcriptomic landscape during mucosal healing. We took advantage of our recent work showing that by day (d)14, the intestinal barrier integrity is restored following damage induced by dextran sodium sulfate (DSS)^10^. Therefore, we treated wild type (WT) mice with DSS in drinking water for 7 days followed by 7 days of recovery and d14 colonic tissue was taken to generate frozen Swiss-rolls to be processed for ST (Fig. 3A). Despite the recovery at a physiological level (i.e. body weight gain) (Extended Data Fig. 4A), the colonic tissue after DSS treatment did not fully return to homeostasis, as demonstrated by its reduced length (a sign of inflammation) (Extended Data Fig. 4B).

**Fig. 3.**
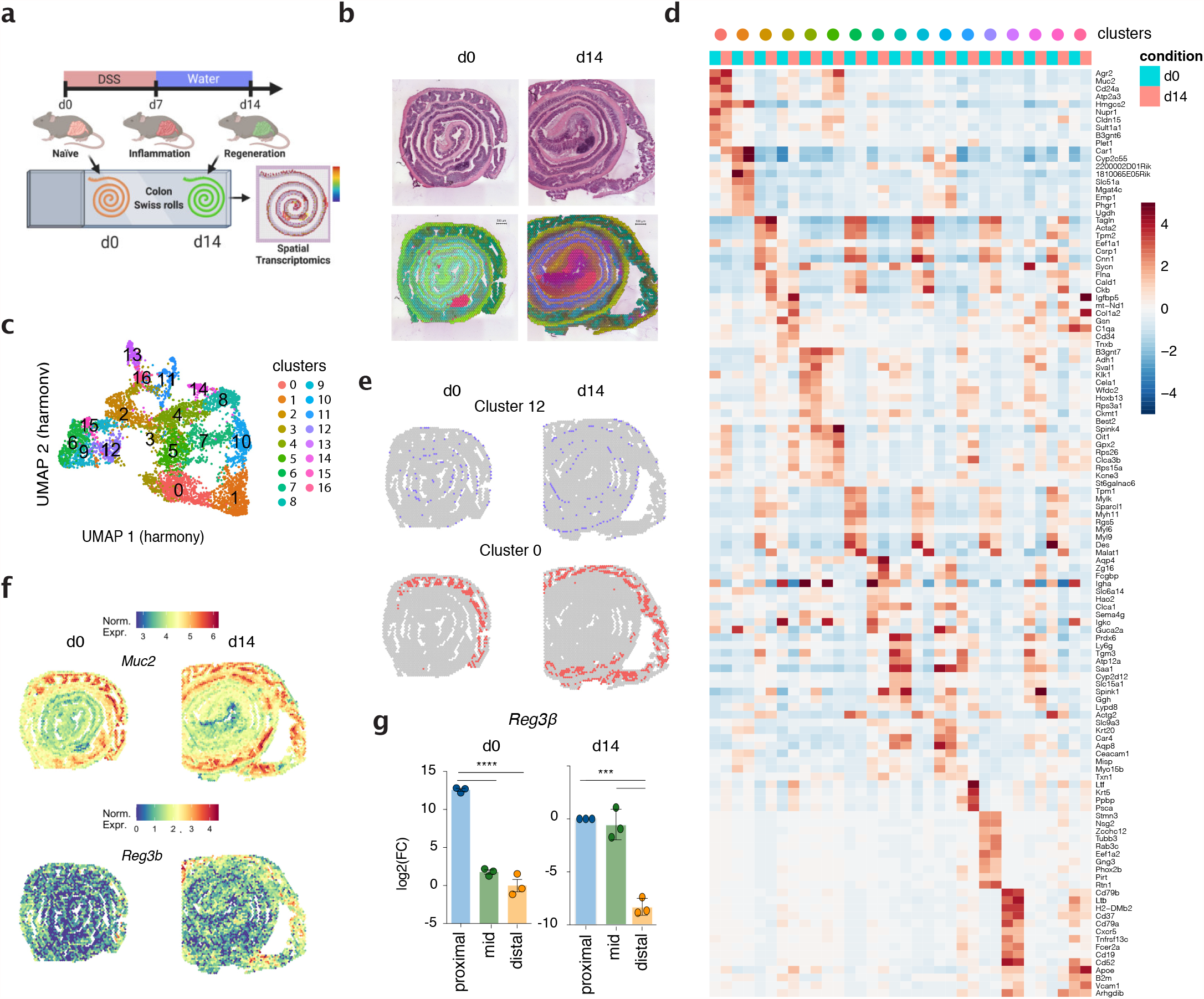
Changes of the molecular topography during mucosal healing are dominant at the distal colon. (A) Schematic representation of the experiment: colitis was induced by dextran sodium sulfate (DSS) administration in drinking water for 7 days followed by 7 days of regular water to promote tissue repair. Colonic tissue from a wild-type naive mouse (d0, from Fig. 1) and from a mouse undergoing colonic regeneration (d14) were processed as Swiss roll for spatial transcriptomic using Visium 10X technology. (B) Top: Hematoxylin and eosin staining of colonic tissue from d0 and d14. Bottom: spatial representation of UMAP values in CMYK colors on colon d0 and d14. Spots with the same color in the two time points represent transcriptionally similar regions. (C) Uniform Manifold Approximation and Projection (UMAP) representation of 16 color-coded clusters defining regional transcriptome diversity in the colonic d0 and d14 datasets combined. (D) Heatmap showing expression of top genes defining each cluster (color-coding on top) in the ST datasets from the two timepoints (light blue columns: colon d0; pink columns: colon d14). (E) Schematic representation of cluster 0 and cluster 12 distribution in colon d0 (on the left) and d14 (on the right). (F) Expression of selected genes in cluster 0 onto ST. (G) qPCR validation of regional expression of *Reg3b* in proximal, mid and distal colonic biopsies from wild type mice at steady state conditions (d0) and during mucosal healing (d14)(n=3, each dot represents one mouse).

At the histological level, large lymphoid patches, as well as the muscle and mucosal layer across the intestine, were easily identified (Extended Data Fig. 4C). Hematoxylin and eosin (H&E) sections annotated by a blinded pathologist revealed the heterogeneity of the tissue, including the presence of isolated lymphoid follicles (ILFs), as well as areas with edema, hyperplasia, crypt duplication, and normal tissue (Extended Data Fig. 4D). Of note, the distal colon (center of the Swiss roll) showed marked alterations, whereas the proximal colon (outer Swiss roll) seemed non-affected (Extended Data Fig. 4D).

The d14 ST dataset consisted of 3630 individual spots with a number of unique genes per spot (nFeature_RNA) that was comparable to the d0 tissue section (Extended Data Fig. 1A). To first appreciate how the process of mucosal healing spatially altered the colonic transcriptome, the ST data from d0 and d14 were embedded in 3 dimensions using Uniform Manifold Approximation and Projection (UMAP). The values of these 3 dimensions were then re-scaled into a unit cube (with a range of 0 to 1) and used as channels in CMYK color space to generate a specific color for each ST spot (Fig. 3B, bottom part). Interestingly, lymphoid follicles (identified by H&E staining) and areas in the proximal colon showed high similarity (i.e. same color) between d0 and d14 samples (Fig. 3C), suggesting that these structures are transcriptionally less affected during the process of mucosal healing following intestinal injury. Vice versa, in line with the histo-pathological scoring, the distal portion of the d14 colon was the most dramatically affected region.

To visualize how the colonic tissue is transcriptionally organized in different areas, we integrated the data from d0 and d14 using harmony ^27^ and performed cluster analysis. We annotated 17 distinct clusters, which were visualized by embedding the data in 2 dimensions with UMAP (Fig. 3C). Differentially up-regulated genes per cluster are summarized in a heatmap showing the top conserved genes in each cluster (Fig. 3D). Each cluster defined a distinct geographic area of the tissue (Extended Data Fig. 5) For instance, cluster 12 designated the ENS, with scattered expression in the submucosal layer, whereas cluster 0 mapped spatially to the proximal colon (Fig. 3E). Interestingly, genes defining cluster 0 expanded towards the mid colon during mucosal healing (d14)(Fig. 3E). Among these genes, *Muc2* and *Reg3b* showing expanded expression towards the mid colon (Fig. 3F) play a key role in establishing the barrier integrity. Using qPCR, we validated the expanded expression of *Reg3b* during mucosal healing (Fig. 3G). Overall, cluster analysis revealed that despite the existence of a conserved transcriptional colonic regionalization, the process of tissue healing underlies the emergence of distinct molecular signatures and alters the distribution of specific gene expression.

### Non-negative matrix factorization analysis revealed a previously unappreciated transcriptomic regionalization during mucosal healing

Even though the majority of the tissue was defined by clusters equally represented on both time points, some clusters displayed a partial or drastic enrichment during tissue healing. Cluster 3 (found in the distal colon at the interface between the LP and the muscularis layer), cluster 11 and 16 (localized in the damaged area of d14 distal colon) and cluster 13 (marking lymphoid follicles) were drastically enriched during d14 (Fig. 4A and S5). To visualize how the process of mucosal healing alters the transcriptomic landscape of the colon, we deconvolved the d0 and d14 datasets jointly into 20 factors using NNMF (Extended Data Fig. 6 and S7). Among these, 8 factors were defined by genes expressed in specific regions during mucosal healing (d14), but not at d0 (Fig. 4B). In the proximal colon, factor 1 was characterized by genes involved in bile acid and fatty acid metabolism (e.g. *Cyp2c55*, an enzyme involved in the metabolism of 19-hydroxyeicosatetranoic acid). By contrast, factors positioned in the distal colon were characterized by genes involved in inflammatory processes (e.g. *Duoxa2* and *Il18*) and tissue remodeling (e.g. *Col1a1* and *Col1a2*), among others (Fig. 4C). Due to their close proximity and their marked enrichment during mucosal healing, we focused on factors 5, 7, 14, and 20. Factor 5 delineated an edematous area, which was histologically characterized by inflammation and positioned right beneath a severely injured epithelial layer with complete loss of crypt architecture (i.e. factor 14). Pathway analysis revealed that factor 5 was associated with processes involving anatomical structure development, cell adhesion, and extracellular matrix (ECM) organization (Fig. 4D). Among the top genes defining this factor, we found *Igfbp5* and *Igfbp4*, as well as collagens *Col1a1* and *Col1a2* (Fig. 4C). In contrast, factor 14 was characterized by the expression of genes involved in stress response (e.g. *Duoxa2* and *Aldh1a3*) and leukocyte infiltration (e.g. *Ly6a* ansd *Cxcl5*), which indicate an acute response to a barrier breach and tissue damage.

**Fig. 4.**
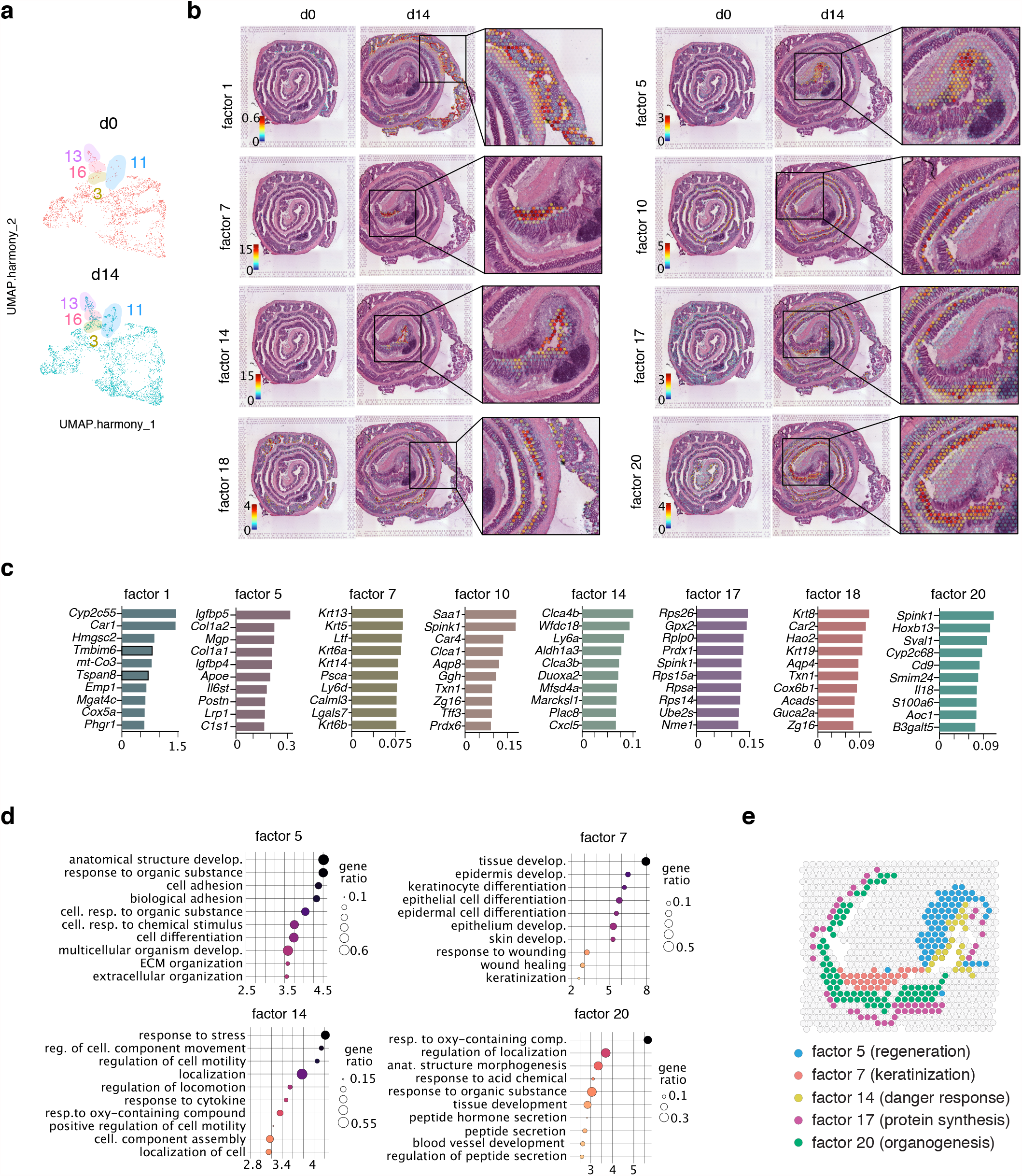
Non-negative matrix factorization reveals eight distinct molecular patterns during colon mucosal healing. (A) UMAP representation of 16 clusters in d0 and d14 colon. (B) Hematoxylin and eosin images displaying overlaid spots with the highest factor weight. (C) Top 10 genes defining the indicated NNMFs (factors). (D) Functional enrichment analysis (Gene Ontology, GO) based on the top genes defining factor 5, 7, 14 and 20. (E) Schematic representation summarizing the expression pattern between selected factors. Biological processes associated with each factor are indicated in brackets.

At the end of the colonic tissue, the anus separates a mucosal tissue with a monolayered epithelium (the rectum) from a stratified squamous epithelium (skin). Homeostatic breakdown resulting from colonic inflammation generates an area of epithelial instability wherein a heterogeneous tissue at the interface between skin and colonic epithelium appears (i.e. enlarged multilayered crypt-like structures with squamous, but not cornified, appearance). Factor 7, characterized by the expression of several keratins (e.g. *Krt13, Krt5, Krt14* and *Krt6a*) and pathways involving keratinocytes differentiation and wound healing, delineated this area (Fig. 4C-D).

Finally, factor 20 predominantly defined the distal epithelium undergoing hyperplasia and crypts arborization, which indicates epithelial repair. Gene ontology revealed that this factor was associated with organogenesis (e.g. *Hoxb13*) and response to organic substrates (e.g. *Cyp2c68*) (Fig. 4C-D). Overall, DSS-induced injury resulted in an assorted co-occurrence of different histopathological processes within the murine colon. Furthermore, our analysis revealed a previously unappreciated heterogeneous transcriptional and regional landscape of tissue repair (Fig. 4E).

### Predictive algorithms revealed coordinated signaling pathways depending on location

We interrogated if distinct signaling pathways could be inferred by the spatially organized transcriptional profiles by using PROGENy ^28,29^. Unlike other Gene Set Enrichment tools (as KEGG), PROGENy estimates signalling pathway activities by looking at expression changes of downstream genes in signaling pathways, which provides a more accurate estimation of the activity of the pathway. A score for each of the 14 pathways annotated in PROGENy (i.e. Wnt, VEGF, Trail, TNFα, TGFβ, PI3K, p53, NFkB, MAPK, JAK/STAT, Hypoxia, Estrogen, Androgen and EGFR) was estimated for each ST spot on d0 and d14 slides (see methods). First, we computed a correlation matrix to understand how the spatially organized transcriptional programs identified by NNMF (i.e. factors of Fig. 4) could be explained by signalling pathway activities. We observed two main groups, in which, group 2 pathways (Androgen, JAK-STAT, NFkB, TNFα, p53, Hypoxia and Trail), characteristic of an inflammatory/acute response to damage, were associated with factors comprising damaged distal epithelium (e.g. factor 7, 10, 14, 20) and proximal epithelium (factor 11, 19)(Fig. 5A-B). In contrast, group 1 pathways (TGFβ, Wnt, PI3K, Estrogen and EGFR), normally regulating pro-regenerative/tissue remodeling processes, were associated with factors defining the tissue beneath the damaged epithelium (e.g. factor 5 and 17), the muscle layer (factor 2, 6 and 12) and lymphoid follicles (factor 9) (Fig. 5B). In addition, MAPK and VEGF pathways were associated with similar factors, and as expected, the spatial patterns of MAPK and VEGF pathways activity were comparable (Fig. 5C). Of note, at steady state conditions (d0) MAPK and VEGF pathways were homogeneously active along the mid-distal colon, whereas during mucosal healing, their activation was higher within the damaged/regenerating areas (Fig. 5C, black arrows).

**Fig. 5.**
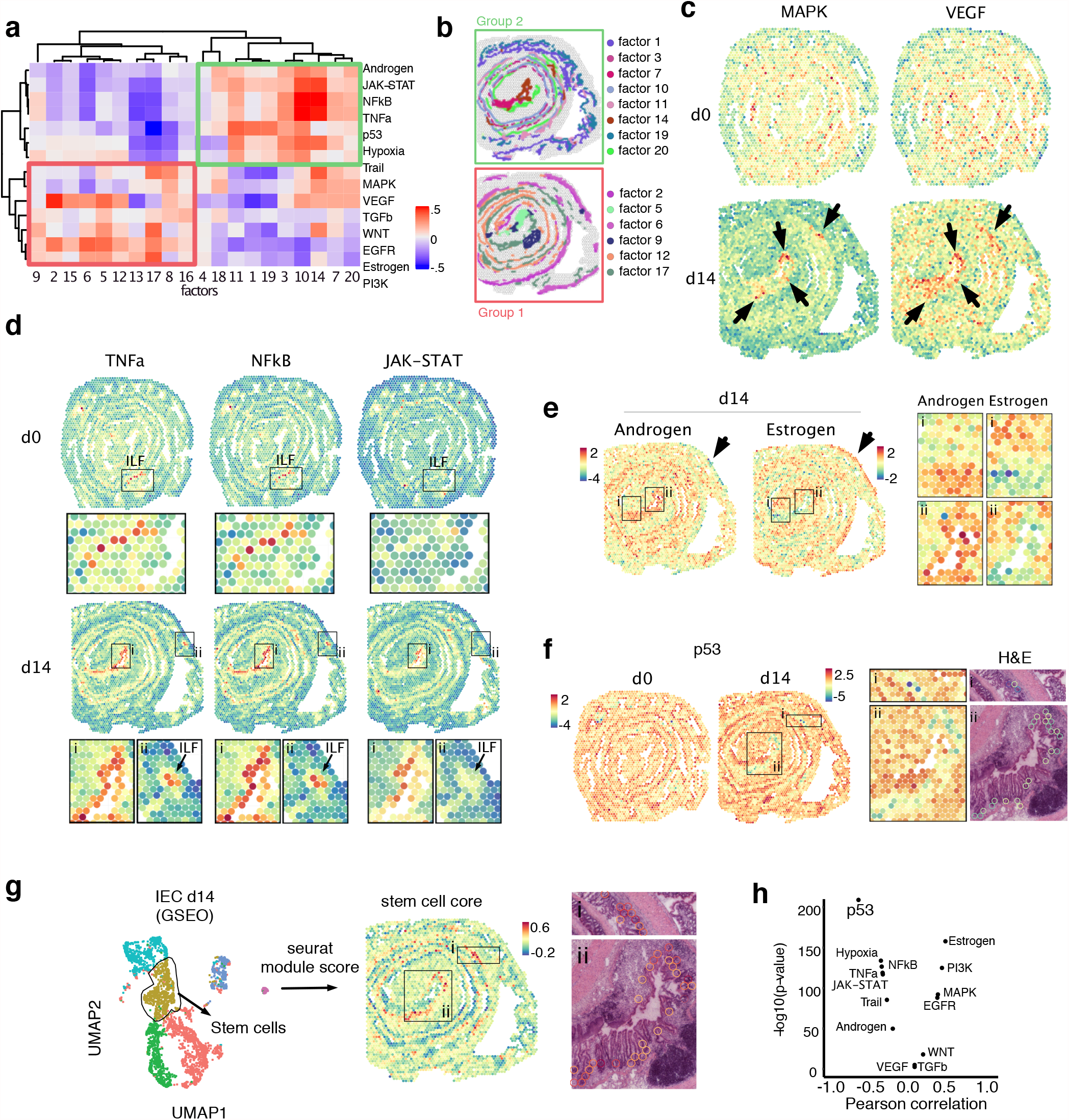
Predictive algorithm reveals pathway-specific spatial patterns during mucosal healing. (A) Correlation matrix between non-negative matrix factorization (NNMF) and pathway activity scores determined by PROGENy (B) Schematic of the colon area at d14 displaying the distribution of the indicated factors. (C) Spatial transcriptomic (ST) spot heatmaps of the colon at d0 (upper swiss rolls) and d14 (lower swiss rolls) showing pathways scores predicted by PROGENy. Arrows indicate areas of tissue damaged as defined by factor 14 shown in Extended Data Fig. 7 and Fig. 4A-C. (D) ST spot heatmaps showing TNFα, NFkB, JAK-STAT pathway activity on d0 (upper Swiss rolls) and d14 (lower Swiss rolls). Selected areas indicated as ILF (isolated lymphoid follicles) or “i” and “ii” and outlined in black are magnified below each Swiss roll. Arrows in “ii” indicate the presence of an ILF as defined by factor 9 Extended Data Fig. 6. (E) Spatial distribution of androgen and estrogen pathway activity at d14. Selected areas (indicated as “i” and “ii”) are magnified. Arrows indicate an example of the muscle layer showing opposite expression patterns between the two pathways. (F) Spatial distribution of p53 pathway activity at d0 and d14. Selected areas indicated as “i” and “ii” on colon d14 are magnified on the right. Hematoxylin and eosin magnifications show the overlaid spots with the lowest p53 activity shown in “i” and “ii”. (G) Left: UMAP visualization of IEC clusters from scRNAseq on colon d14 (Frede et al, unpublished; GSE: 163638). Middle: ST spots from colon d14 are color-coded based on the enrichment of stem cell core signature identified from scRNAseq dataset. Selected areas indicated as “i” and “ii” on colon d14 are magnified on the right. Right: Hematoxylin and eosin magnifications showing the overlaid spots with the highest stem cell signature shown in “i” and “ii”. (H) Pearson correlation between PROGENy predicted pathways and stem cell signature on the ST dataset.

### Shared and complementary pathway activities during mucosal healing

Comparable TNFα, NFkB, and JAK-STAT pathway activation scores between some factors (e.g. factor 10 and 14) (Fig. 5A) suggest interconnectivity between these inflammatory pathways. To test this possibility we further analyze the spatial pattern of these pathways. In particular, the activities of the TNFα and NFkB pathways were almost identical within the colon, regardless of the time point analyzed (Fig. 5D). Higher TNFα and NFkB activities were appreciated in areas associated with injury and ILFs (Fig. 5D). Of note, in the absence of damage/inflammation (d0), the spatial distribution of TNFα and NFkB showed activity confined to the ILF luminal edge (Fig. 5D, d0), in agreement with previous studies showing that TNF drives ILF organogenesis ^30^. Whether ILF forms where subclinical local damage occurs or whether their presence, which allows dynamic exchange with the external environment, causes subclinical inflammation, remains to be explored.

In contrast, JAK-STAT pathway activation showed co-occurrence with TNF and NFkB mostly in the damaged area (factor 14), but not in ILFs (Fig. 5D). These results suggest that although all three pathways may play a role within the damaged tissue, TNFα and NFkB, but not JAK-STAT, are involved in the formation/function of ILFs. Unlike these pathways, androgen and estrogen pathways showed mutually exclusive patterns of activity. Higher androgen pathway activity was observed in areas of injured epithelium, while higher estrogen activity was associated with the muscle layer (arrows, Fig. 5E), suggesting that these pathways negatively regulate each other during mucosal healing.

### Low p53 pathway activity is associated with proliferating crypts

Activation of the p53 pathway was homogeneously distributed across the proximal-distal axis, but it showed more activity in the luminal side compared with the LP and muscle layer (Fig. 5F). Interestingly, p53 activity was lower in the damaged area (box ii in Fig. 5F). Activation of p53 triggers cell cycle arrest, senescence, and apoptosis ^31^, suggesting that spots with decreased p53 activity might be enriched in proliferating cells within the damaged area. To test this possibility, we overimposed lower p53 activity spots onto the H&E images and showed co-localization with the bottom of crypts (Fig. 5F, H&E boxes). To investigate if proliferating stem cell signatures co-localize with low p53 activity spots, we took advantage of our single cell RNA sequencing (scRNAseq) dataset of intestinal epithelial cells from d14 colon and identified a population of proliferating stem cells (Fig. 5G). We mapped the stem cell core signature onto the d14 colon ST datasets, and we superimposed the spots with high scores in the stem cell core onto the H&E section. In agreement with our hypothesis, spots containing high scores (Fig. 5G) coincided with low p53 activity (Fig. 5F). Pearson correlation analysis confirmed that ST spots with high stem cell scores negatively correlated with p53 activity (Fig. 5H). Thus, our data suggest that low p53 activity allows the identification of proliferating crypts during mucosal healing. In summary, we spatially positioned clinically relevant pathways predicted by PROGENy and showed that these pathways are highly coordinated during mucosal healing.

### Integration of human datasets with mouse spatial transcriptomic

To enquire about the translational potential of the murine colonic ST, we investigated whether human datasets could be integrated into murine ST data. Towards this end, we took advantage of a human developing gut dataset ^32^ and mapped 31 distinct epithelial and stromal cells onto our ST dataset (Fig. 6A). We observed correlations between human cells types and distinct murine ST factors (Fig. 6B), indicating specific localization of human cell signatures within the mouse colon. Interestingly, the signature of human proximal enterocytes is highly correlated with factor 1 (Fig. 6B), defining the most proximal epithelium in mice (Extended Data Fig. 8A). Human distal enterocytes and absorptive cells highly correlated with factors 3 and 10 (Fig. 6B), which defines the most distal epithelium in mice (Extended Data Fig. 8A). These results indicate that the transcriptomic features defining proximal and distal epithelial cells are conserved between mouse and humans. On the other hand, two human stromal cells characterized by the expression of the chemokines CCL21 and CXCL13 uniquely and strongly correlated with factor 9, defining lymphoid follicles (Fig. 6B), which is in agreement with the well-known role of these chemokines in ILF development ^33^. Interestingly, these stromal cells mapped in complementary patterns within the mouse ILF (Fig. 6C), suggesting that the coordinated action of these cells might determine the recruitment/localization of immune cells within the follicle. Next, we analyzed the damage/regeneration area (factor 5 and 14) which correlated with S1 (Stromal 1, fibroblast marking bulk of submucosal structural cells in human), S1-COL6A5 and S1-IFIT3 human cells (two subtypes of S1) (Fig. 6B) and mapped in a complementary pattern of distribution (Extended Data Fig. 8B).

**Fig. 6.**
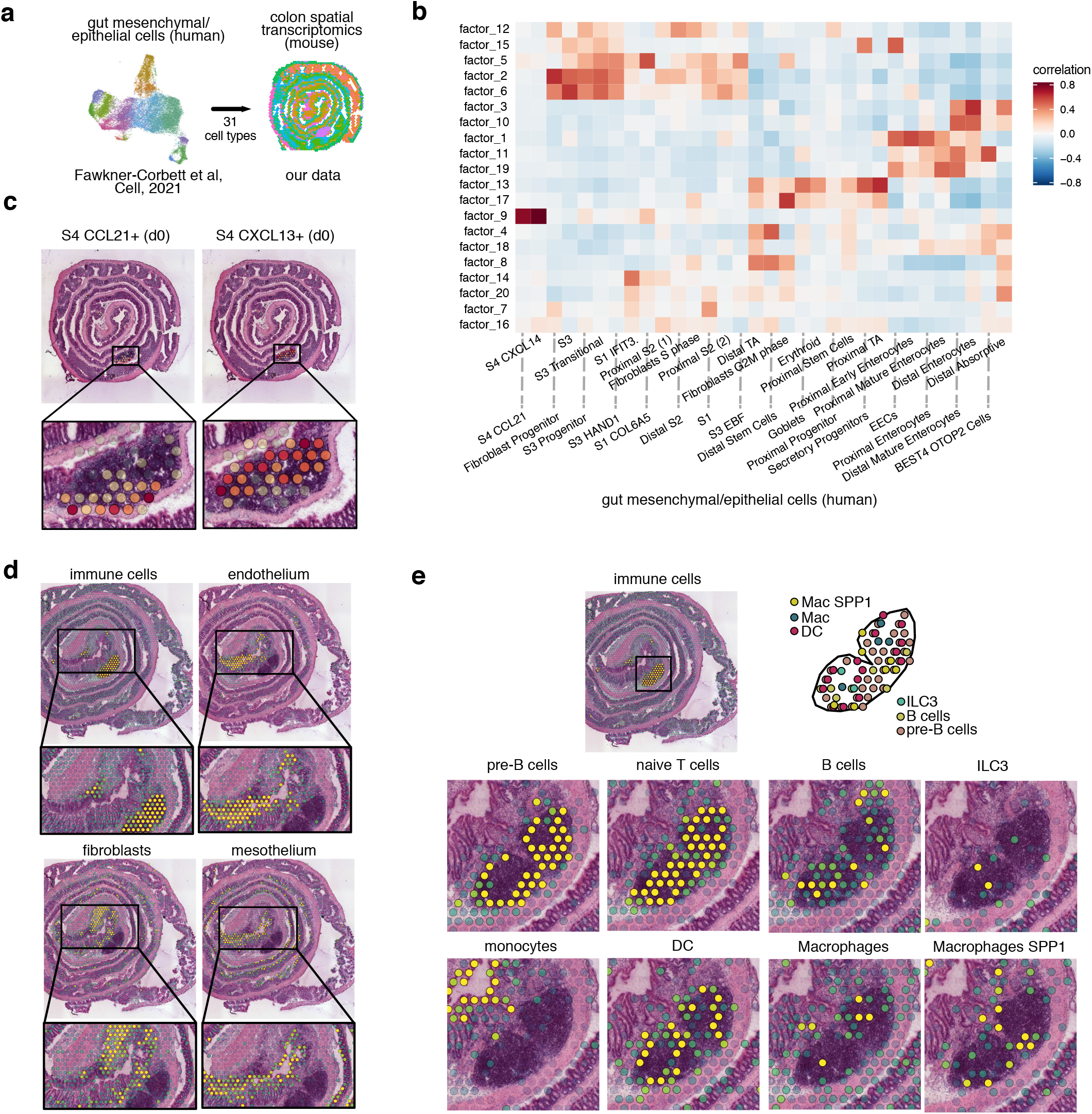
Human cell type mapping onto murine spatial transcriptomic datasets. (A) Scheme showing the integration of published human single cell RNAseq ^32^ and our mouse Visium datasets. (B) Correlation matrix between transcriptomic profiles from human single cell datasets ^32^ and factors defining transcriptomics patterns in mouse ST. (C) Integration of human stromal cell transcriptomic profiles (S4.CCL21+ and S4.CXCL13+) onto visium datasets at day 0. (D) Integration of human intestinal cell transcriptomic profiles onto visium datasets at day 14. (E) Integration of human immune cell transcriptomic profiles onto visium datasets at day 14 and magnification of the isolated lymphoid follicle area.

We then extend our analysis to other cell types during mucosal healing (Fig. 6D and Extended Data Fig. 8C). Interestingly, immune cells, mesothelium, endothelium and fibroblast signatures were spatially enriched within defined areas during mucosal healing. In particular, fibroblasts were dominant in factor 5 (remodeling) and immune cells in factor 9 (lymphoid follicles), whereas endothelial and mesothelial cells colocalized within factor 7 (keratinization) (Fig. 6D and Extended Data Fig. 8D). Among 10 distinct immune cell types identified ^32^, monocytes and SPP1+ macrophages were enriched in factor 14 (danger response) and in factor 5 (tissue remodeling) respectively, in line with their known roles in acute response to injury and matrix deposition/wound healing (Extended Data Fig. 8E). Lymphocytes and dendritic cells, instead, were enriched in factor 9 (lymphoid follicles) (Fig. 6E and Extended Data Fig. 8E). Further analysis showed how these immune cells are heterogeneously distributed within the ILF (Fig. 6E). These results provide a proof-of-concept and support the notion that principles of spatial distribution within the colonic tissue appear to be conserved between species and highlight murine ST as a valuable platform for exploring and translating findings on distribution patterns of cells/genes within a tissue.

### Mapping transcriptomic datasets onto ST to inform medical practice

In order to establish a framework to integrate existing knowledge with ST datasets, we took advantage of our longitudinal RNAseq dataset of colonic tissue collected during acute epithelial injury and the recovery phase in the DSS-induced colitis model ^10^. In this study, we identified sets of genes (called modules) displaying characteristic expression patterns, with some genes being: a) downregulated upon injury (modules 2, 8, 7), b) upregulated during the acute/inflammatory phase (modules 1, 3, 4, 9), and c) upregulated during the recovery phase of DSS colitis (modules 5, 6) ^10^. To identify whether the different temporally regulated processes (i.e. modules) were enriched in specific areas of the tissue, we computed a correlation matrix between the gene signature of modules and ST factors (Fig. 7A). Higher gene enrichment was found in module (m)1 and m6, characterized by genes induced during the inflammatory and recovery phase, respectively (Fig 7A). Among the genes shared between factor 9 and m1, *Ptprc, Cd72* and *Lyz2* encode for proteins expressed by immune cells and map predominantly to lymphoid follicles and damaged areas in d14 (Fig. 7Bi). In contrast, ∼60% of the top driving genes defining factor 15 (i.e. ENS) were shared with m6, and their expression was distributed in the submucosa/muscularis layer where neuronal bodies reside (Fig. 7Ci). GO enrichment analysis of m1 and m2 also confirmed that the most dominant pathways were involved with inflammatory responses and chemical synaptic transmission (Fig. 7Bii and 7Cii). The correlation between temporal transcriptomic modules and spatial factors suggests that factor 9 (lymphoid follicles) and factor 15 (ENS) are characterized by an ongoing inflammatory and regenerative profile, respectively. Thus, the integration of longitudinal and ST data can be a powerful tool to unveil biological processes related to diseases in time and space.

**Fig. 7.**
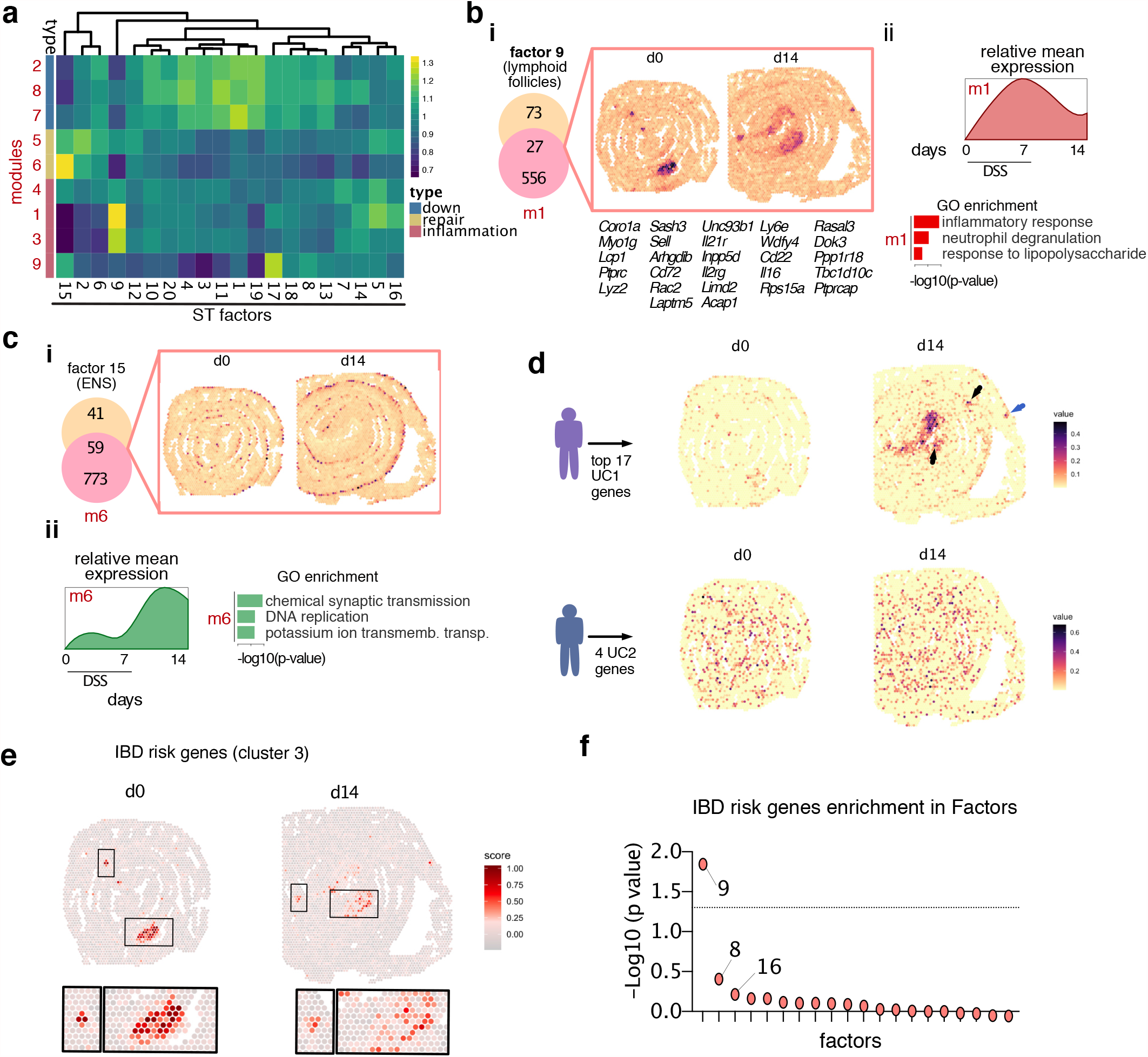
Spatial transcriptomic (ST) allows mapping transcriptomic signatures with clinical relevance. (A) Correlation matrix between transcriptomic modules distinguishing the processes of inflammation and mucosal healing during DSS-induced colitis ^10^ and factors defining transcriptomics patterns in ST. (B) (i) Venn diagram and spatial representation of overlapping genes between module 1 and factor 9. (ii) relative mean expression and Gene Ontology (GO) of genes belonging to module 1. (C) (i) Venn diagram and spatial representation of overlapping genes between module 6 and factor 15. (ii) relative mean expression and Gene Ontology (GO) of genes belonging to module 6. (D) Spatial distribution of genes defining UC1 and UC2 patients on mouse ST colon d0 and d14. (E) Spatial distribution of IBD risk genes from Cluster 3 (Extended Data Fig. 9B) on colon d0 (left) and d14 (right). (F) Gene Set Enrichment Analysis for IBD risk genes in Cluster 3 (Extended Data Fig. 9B) and NNMF factors (Extended Data Fig. 6-7).

### Spatial distribution of genes defining UC1 and UC2 profiles

We then sought to investigate if clinically relevant patient gene signatures could be mapped onto mouse ST datasets. Toward this, we used the recent gene signature identifying ulcerative colitis (UC) subgroups of patients: UC1 and UC2 ^10^. This molecular classification is clinically relevant because the UC1-related transcriptomic signature is associated with poor responses to biological therapies and is enriched with genes involved in neutrophil activity. In contrast, approximately 70% of UC2 patients achieved a clinical response to *anti-TNF* antibodies ^10^. In addition, Smillie et al., ^34^ showed that inflammatory fibroblasts and monocytes mainly drive anti-TNF resistance, and many of the genes upregulated in UC1 patients colonic tissue are highly expressed by these cell types. To further understand the spatial distribution of differentially expressed genes between UC1 and UC2 patients, we mapped all genes upregulated in either UC1 and UC2 patients onto colon ST. Whereas genes defining UC2 patients were homogeneously expressed across the colon (d0 and d14) (Fig. 7D), genes defining UC1 were mostly localized within the damage/repair area at d14 (Fig. 7D). Further investigation revealed that all the main functional classes upregulated in UC1 patients, including collagen synthesis (*Col12a1, Col4a1, Col4a2, Col7a1*), ECM breakdown (*Mmp3*), Wnt-signalling pathway (*Wnt5a*), cytokine signalling (*Il11, Il1b, Il33, Il1r2, Il1rn, Tnfrsf11b, Csf2rb, Csf3r, Socs3, Trem1, Cxcr2*) and innate immunity (*S100a9, S100a8, C5ar1, Sell, S100a4*), were also upregulated in areas of tissue damage Fig. S9A). In summary, these results suggest that UC1 patients may possess higher tissue damage and ulceration compared with UC2 patients.

### Defining the topography of IBD risk genes

Spatial transcriptomics of the colon undergoing injury/repair provides an opportunity to comprehensively map IBD-associated risk genes. Therefore, we interrogated the expression pattern of various human IBD-risk genes ^35-39^ on the ST profile of d0 and d14 colonic tissue. Out of the 122 interrogated genes, 95 IBD-risk genes were selected based on the existence of their murine ortholog and their detectable expression in the ST dataset. In order to identify whether the spatial expression of these variants defined topographic patterns within the tissue, we computed a correlation matrix (Extended Data Fig. 9B). Cluster analysis resulted in three main co-expression clusters, with cluster 3 possessing the highest spatial expression correlation between genes (Extended Data Fig. 9B). Functional annotation of the genes within this cluster revealed enrichment in pathways related to immune cell recruitment (e.g. *Itgal, Icam1, Itga4*), activation (e.g. *Cd6, Plcg2, Ncf4, Il10ra*), and antigen presentation (e.g. *Tap1, Tap2, Psmb8*).

To understand the spatial distribution of genes belonging to cluster 3, we mapped them onto the colonic tissue using ModuleScore, a Seurat function assigning a score in the ST dataset to a set of predefined genes (i.e. cluster 3 genes). In line with the functional annotation of cluster 3 genes, we observed enrichment in lymphoid follicles areas both on d0 and d14 (Fig. 7E). To understand which ST factors were enriched with IBD risk genes, we performed Gene Set Enrichment Analysis (GSEA) and calculated overrepresentation scores of IBD-risk variants in the NNMF dataset (Extended Data Fig. 6-7). This analysis showed that factor 9, defining lymphoid follicles within the tissue (Extended Data Fig. 6), was the only factor with significant enrichment of IBD risk genes (Fig. 7F). Altogether, this analysis revealed that the expression of a subset of human IBD-risk genes spatially co-occur within the murine colon. Their specific expression pattern suggests that colonic tissue lymphoid follicles might define the area to potentially target when developing therapeutic strategies for IBD patients displaying aberrant immune activation.

## Discussion

We and others have deeply characterized the transcriptomic landscape during mucosal healing in the colon and small bowel ^10,40,41^. However, these studies lacked the spatial resolution describing where genes were expressed. Here, we spatially placed cell populations and pathways that might play pivotal roles in driving tissue response to damage. The current study uncovered spatial transcriptomic patterns that are present at steady state conditions and that arise in response to damage; these spatial transcriptomic patterns were characterized by unique transcriptional signatures and coincided with different histological processes. Moreover, we profiled the regional distribution of different biological processes, such as acute response to injury or a regenerative response. Finally, we demonstrated the clinical relevance of this dataset as seen by conserved spatial localization of gene signatures in human tissue and transcriptomics data and by testing the distribution of clinically-relevant genes.

A recent study characterized the transcriptomic landscape during human gut development ^32^. Here, we further these results by taking advantage of murine colonic Swiss rolls that fit the 6.5 mm^2^ area constraints provided by the manufacturer to perform spatial transcriptomics. This approach enabled us to visualize the transcriptomic landscape of the whole colon in the same slide, including the most proximal and distal segments. In combination with bioinformatics tools (NNFM analysis), we uncovered a previously unappreciated molecular regionalization of the colonic tissue in steady state conditions. This analysis allowed the identification of distinct epithelial, LP, and muscularis/submucosa genetic programs depending on their proximal to distal colon localization. Importantly, when mapping human cells into our ST datasets, we observed conservation in transcriptomic features defining proximal and distal locations, suggesting that our newly described molecular segmentation is conserved across mammals.

Using the entire murine colon, we provided a detailed analysis of a previously unappreciated compartmentalization of the tissue repair process. In line with previous reports ^42^, our unbiased analysis of the transcriptomic landscape during mucosal healing reveals that while dramatic transcriptomic changes occur in the distal colon, the proximal colon remains almost comparable to the steady state. Two potential scenarios can be proposed: a) the level of damage is homogenous and the proximal colon heals faster compared with the distal colon; or b) the proximal colon is more protected compared with the distal colon. Dramatic changes in the distal rather than the proximal colon are in agreement with the phenotype observed in UC patients, where the focus of inflammation extends proximally from the rectum ^43^. In addition, genes and/or pathways, such as the JAK-STAT and TNFα pathway ^44^ or genes/pathways characterizing UC1 patients ^10^, were found to be dominant in the distal colon, suggesting that DSS-induced colitis is a clinically relevant experimental model of UC1. Whether higher levels of damage/tissue repair in the distal colon depend on microbiota, host-induced responses to the microenvironment, or just different kinetics, remains to be addressed.

At steady state conditions, we identified molecular signatures associated with lymphoid structures. Unlike Peyer’s patches (PP) that are macroscopically visible in the murine small intestine, CP and ILF cannot be dissected and analyzed separately from the rest of the colonic tissue for transcriptomic readouts. For instance, enrichment in the NFkB and TNFα pathway activity was detected in the lumen-facing area corresponding to the epithelial layer overlaying lymphoid clusters in steady state. Because these pathways are usually associated with immune activation/inflammatory responses, these data suggest that ILF-associated epithelium is undergoing inflammation. Moreover, expression of clusterin (*Clu*) alone was found to be highly specific and sufficient in defining the isolated lymphoid follicles (ILF). Previous studies have reported Clu expression in follicular dendritic cells (FDC), ^45^ as well as M cells ^21^ in the Peyer’s patches. While *Clu* expression in FDCs serves as a pro-survival factor for germinal center B cells in the follicle, the role of Clu in M cells is not clear. Interestingly, a recent study ^40^ identified *Clu* as a marker of intestinal stem cells (ISC), known as revival stem cells, which are rarely found in steady state, but are predominantly found in regenerating intestine following injury. Whether the expression of *Clu* in the ILF- and follicle-associated epithelium has any bearing during colonic infection and regeneration needs to be further investigated.

Besides the proximal-distal variance in the transcriptomic alterations during tissue repair, our NNMF analysis revealed a high degree of compartmentalization within the distal colon itself. At least 5 factors were delineating topographically and transcriptionally distinct areas in the distal colon undergoing different biological responses to tissue injury. Such heterogeneity is likely the result of different healing programs, such as skin-like re-epithelialization (factor 7), acute damage (factor 14) and tissue regeneration (factor 20). Supporting this notion, the integration of our RNAseq kinetic dataset ^10^ into the ST map revealed that certain areas of the distal colon were transcriptionally closer to samples from the acute phase of DSS colitis (i.e. d6 to d8). The asynchronous nature of the healing process may be associated with varying degrees of exposure to the external environment and the elaborate architecture of the tissue. Similarly, in human IBD, the “patchiness of the inflammatory response” is a well-known characteristic of Crohn’s disease. In UC, the inflammation is traditionally thought to be continuous with increasing intensity in distal colon, but longitudinal sampling has revealed episodes of both macroscopic and microscopic patchiness of inflammation ^46^. Our dataset thus provides a valuable resource to interrogate the transcriptional programs underlying distinct temporal and biological processes of tissue healing.

PROGENy allows the prediction of pathways activated in specific regions of the colon. Our analysis revealed a strong correlation between pathway activities, such as TNFα and NFkB, suggesting that one pathway might depend completely on the other. We also identified that the area of injury is characterized by several pathways that are also increased on the ILF edge at steady state conditions, suggesting that the formation of lymphoid follicles result in the induction of damage-associated pathways. Finally, decreased p53 activity represents a good strategy to identify damage-associated proliferating crypts. These data suggest that within the same region, intestinal crypts are heterogeneous in their response to damage; our dataset provides a toolkit to investigate this composite response.

### ST to inform clinical practice

Our study provides evidence that ST can be used to map clinically relevant genes and pathways. Genes characterizing a newly described UC subgroup were associated with poor treatment response in the damaged area of the regenerating colon. Previous studies confirm these results; increased inflammation severity predicted poor response to anti-TNF treatments ^47^ or blocking cell recruitment to the inflamed intestine using anti-a4b7 antibodies ^48^. As a result of severe inflammation, colonic ulceration may lead to lower therapeutic responses due to decreased blood drug concentration/drug leakage ^49^. Our data suggest that UC1 patients have increased tissue damage compared with UC2 patients, which might contribute to their poor response to biological therapies.

## Acknowledgments

We thank members of the Villablanca lab for helpful comments and Maria Lim for editorial assistance. We thank Kim Thrane for her invaluable expertise in setting up the Visium experiments and for help with tissue imaging. C.E. was supported by the Marie Sklodowska-Curie grant agreement No 844712. L.L. was funded by grants from the Helmsley foundation. E.J.V. was supported by grants from the Swedish Research Council, VR grant K2015-68X-22765-01-6 and 2018-02533, Formas grant nr. FR-2016/0005, Cancerfonden (19 0395 Pj), and the Wallenberg Academy Fellow program (2019.0315). The computations and data handling were enabled by resources provided by the Swedish National Infrastructure for Computing (SNIC) at KTH partially funded by the Swedish Research Council through grant agreement no. 2018-05973. Some schematics were partially created with BioRender.com.

## Author Contributions

SMP, SD, AF, OED, RM, XL, GM, and CE performed experiments. LL, ROR, KPT, and KS performed bioinformatics analysis of the transcriptomic data. EJV and SD conceived the idea. NG, JSR, and JL provided resources. SMP, LL, and EJV wrote the paper. All authors discussed the data, read, and approved the manuscript.

## References

1. Li, N., et al. Spatial heterogeneity of bacterial colonization across different gut segments following inter-species microbiota transplantation. Microbiome 8, 161 (2020).

2. Mowat, A.M. & Agace, W.W. Regional specialization within the intestinal immune system. Nat Rev Immunol 14, 667–685 (2014).

3. Fenton, T.M., et al. Immune Profiling of Human Gut-Associated Lymphoid Tissue Identifies a Role for Isolated Lymphoid Follicles in Priming of Region-Specific Immunity. Immunity 52, 557–570 e556 (2020).

4. Villablanca, E.J., et al. MyD88 and retinoic acid signaling pathways interact to modulate gastrointestinal activities of dendritic cells. Gastroenterology 141, 176–185 (2011).

5. Guan, Q. A Comprehensive Review and Update on the Pathogenesis of Inflammatory Bowel Disease. J Immunol Res 2019, 7247238 (2019).

6. Blanpain, C., Horsley, V. & Fuchs, E. Epithelial stem cells: turning over new leaves. Cell 128, 445–458 (2007).

7. Santos, A.J.M., Lo, Y.H., Mah, A.T. & Kuo, C.J. The Intestinal Stem Cell Niche: Homeostasis and Adaptations. Trends Cell Biol 28, 1062–1078 (2018).

8. Brazil, J.C., Quiros, M., Nusrat, A. & Parkos, C.A. Innate immune cell-epithelial crosstalk during wound repair. J Clin Invest 129, 2983–2993 (2019).

9. Owens, B.M. & Simmons, A. Intestinal stromal cells in mucosal immunity and homeostasis. Mucosal Immunol 6, 224–234 (2013).

10. Czarnewski, P., et al. Conserved transcriptomic profile between mouse and human colitis allows unsupervised patient stratification. Nat Commun 10, 2892 (2019).

11. Stahl, P.L., et al. Visualization and analysis of gene expression in tissue sections by spatial transcriptomics. Science 353, 78–82 (2016).

12. Lin, X. & Boutros, P.C. Optimization and expansion of non-negative matrix factorization. BMC Bioinformatics 21, 7 (2020).

13. Uhlen, M., et al. Proteomics. Tissue-based map of the human proteome. Science 347, 1260419 (2015).

14. Baumgartner, W. Possible roles of LI-Cadherin in the formation and maintenance of the intestinal epithelial barrier. Tissue Barriers 1, e23815 (2013).

15. Tetteh, P.W., et al. Generation of an inducible colon-specific Cre enzyme mouse line for colon cancer research. Proc Natl Acad Sci U S A 113, 11859–11864 (2016).

16. Triff, K., et al. Genome-wide analysis of the rat colon reveals proximal-distal differences in histone modifications and proto-oncogene expression. Physiol Genomics 45, 1229–1243 (2013).

17. Tan, C.W., Hirokawa, Y., Gardiner, B.S., Smith, D.W. & Burgess, A.W. Colon cryptogenesis: asymmetric budding. PLoS One 8, e78519 (2013).

18. Brennan, F.E. & Fuller, P.J. Acute regulation by corticosteroids of channel-inducing factor gene messenger ribonucleic acid in the distal colon. Endocrinology 140, 1213–1218 (1999).

19. Renes, I.B., et al. Alterations in Muc2 biosynthesis and secretion during dextran sulfate sodium-induced colitis. Am J Physiol Gastrointest Liver Physiol 282, G382–389 (2002).

20. Fleming, R.E., et al. Carbonic anhydrase IV expression in rat and human gastrointestinal tract regional, cellular, and subcellular localization. J Clin Invest 96, 2907–2913 (1995).

21. Verbrugghe, P., Kujala, P., Waelput, W., Peters, P.J. & Cuvelier, C.A. Clusterin in human gut-associated lymphoid tissue, tonsils, and adenoids: localization to M cells and follicular dendritic cells. Histochem Cell Biol 129, 311–320 (2008).

22. Castro, C.D. & Flajnik, M.F. Putting J chain back on the map: how might its expression define plasma cell development? J Immunol 193, 3248–3255 (2014).

23. He, H., et al. CCR6(+) B lymphocytes responding to tumor cell-derived CCL20 support hepatocellular carcinoma progression via enhancing angiogenesis. Am J Cancer Res 7, 1151–1163 (2017).

24. Suan, D., et al. CCR6 Defines Memory B Cell Precursors in Mouse and Human Germinal Centers, Revealing Light-Zone Location and Predominant Low Antigen Affinity. Immunity 47, 1142–1153 e1144 (2017).

25. Savage, A.K., Liang, H.E. & Locksley, R.M. The Development of Steady-State Activation Hubs between Adult LTi ILC3s and Primed Macrophages in Small Intestine. J Immunol 199, 1912–1922 (2017).

26. Reinicke, A.T., et al. Ubiquitin C-terminal hydrolase L1 (UCH-L1) loss causes neurodegeneration by altering protein turnover in the first postnatal weeks. Proc Natl Acad Sci U S A 116, 7963–7972 (2019).

27. Korsunsky, I., et al. Fast, sensitive and accurate integration of single-cell data with Harmony. Nat Methods 16, 1289–1296 (2019).

28. Schubert, M., et al. Perturbation-response genes reveal signaling footprints in cancer gene expression. Nat Commun 9, 20 (2018).

29. Holland, C.H., Szalai, B. & Saez-Rodriguez, J. Transfer of regulatory knowledge from human to mouse for functional genomics analysis. Biochim Biophys Acta Gene Regul Mech 1863, 194431 (2020).

30. Furtado, G.C., et al. TNFalpha-dependent development of lymphoid tissue in the absence of RORgammat(+) lymphoid tissue inducer cells. Mucosal Immunol 7, 602–614 (2014).

31. Hafner, A., Bulyk, M.L., Jambhekar, A. & Lahav, G. The multiple mechanisms that regulate p53 activity and cell fate. Nat Rev Mol Cell Biol 20, 199–210 (2019).

32. Fawkner-Corbett, D., et al. Spatiotemporal analysis of human intestinal development at single-cell resolution. Cell 184, 810–826 e823 (2021).

33. Mebius, R.E. Organogenesis of lymphoid tissues. Nat Rev Immunol 3, 292–303 (2003).

34. Smillie, C.S., et al. Intra- and Inter-cellular Rewiring of the Human Colon during Ulcerative Colitis. Cell 178, 714–730 e722 (2019).

35. de Lange, K.M., et al. Genome-wide association study implicates immune activation of multiple integrin genes in inflammatory bowel disease. Nat Genet 49, 256–261 (2017).

36. Uniken Venema, W.T., Voskuil, M.D., Dijkstra, G., Weersma, R.K. & Festen, E.A. The genetic background of inflammatory bowel disease: from correlation to causality. J Pathol 241, 146–158 (2017).

37. Verstockt, B., Smith, K.G. & Lee, J.C. Genome-wide association studies in Crohn’s disease: Past, present and future. Clin Transl Immunology 7, e1001 (2018).

38. Liu, J.Z. & Anderson, C.A. Genetic studies of Crohn’s disease: past, present and future. Best Pract Res Clin Gastroenterol 28, 373–386 (2014).

39. Jostins, L., et al. Host-microbe interactions have shaped the genetic architecture of inflammatory bowel disease. Nature 491, 119–124 (2012).

40. Ayyaz, A., et al. Single-cell transcriptomes of the regenerating intestine reveal a revival stem cell. Nature 569, 121–125 (2019).

41. Lukonin, I., et al. Phenotypic landscape of intestinal organoid regeneration. Nature 586, 275–280 (2020).

42. Yan, Y., et al. Temporal and spatial analysis of clinical and molecular parameters in dextran sodium sulfate induced colitis. PLoS One 4, e6073 (2009).

43. Tontini, G.E., Vecchi, M., Pastorelli, L., Neurath, M.F. & Neumann, H. Differential diagnosis in inflammatory bowel disease colitis: state of the art and future perspectives. World J Gastroenterol 21, 21–46 (2015).

44. Troncone, E., Marafini, I., Del Vecchio Blanco, G., Di Grazia, A. & Monteleone, G. Novel Therapeutic Options for People with Ulcerative Colitis: An Update on Recent Developments with Janus Kinase (JAK) Inhibitors. Clin Exp Gastroenterol 13, 131–139 (2020).

45. Huber, C., et al. Lymphotoxin-beta receptor-dependent genes in lymph node and follicular dendritic cell transcriptomes. J Immunol 174, 5526–5536 (2005).

46. Kim, B., Barnett, J.L., Kleer, C.G. & Appelman, H.D. Endoscopic and histological patchiness in treated ulcerative colitis. Am J Gastroenterol 94, 3258–3262 (1999).

47. Arias, M.T., et al. A panel to predict long-term outcome of infliximab therapy for patients with ulcerative colitis. Clin Gastroenterol Hepatol 13, 531–538 (2015).

48. Barre, A., Colombel, J.F. & Ungaro, R. Review article: predictors of response to vedolizumab and ustekinumab in inflammatory bowel disease. Aliment Pharmacol Ther 47, 896–905 (2018).

49. Brandse, J.F., et al. Loss of Infliximab Into Feces Is Associated With Lack of Response to Therapy in Patients With Severe Ulcerative Colitis. Gastroenterology 149, 350–355 e352 (2015).

