## Supplementary material for "The spatial transcriptomic landscape of the healing intestine following damage": supp figures

### Extended data Fig. 1

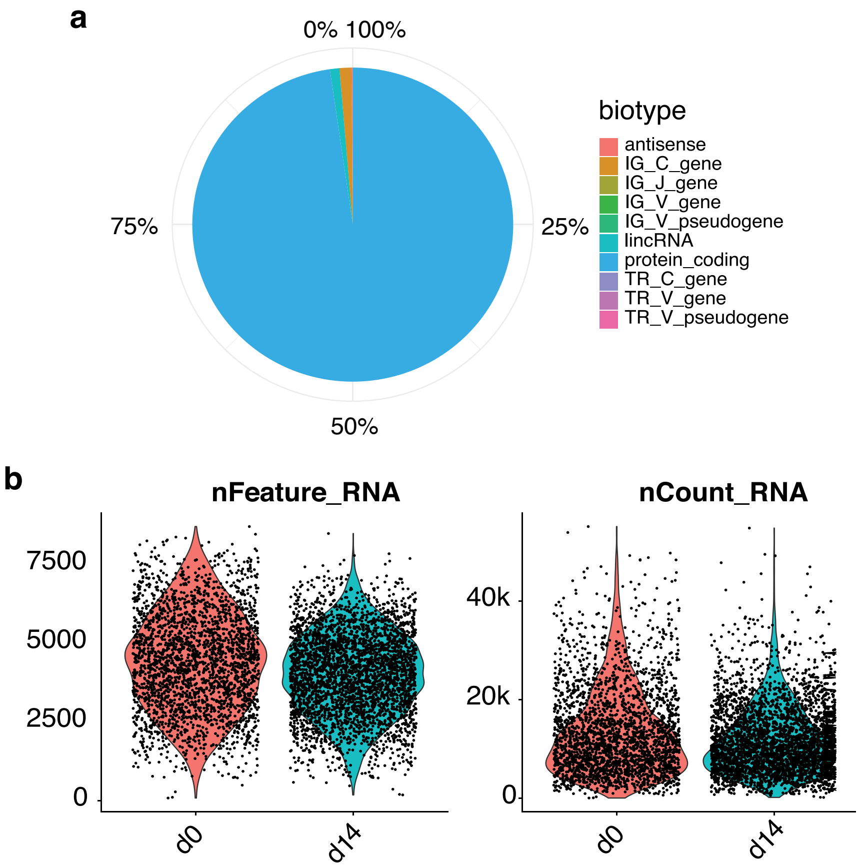

### Extended data Fig. 2

**a**

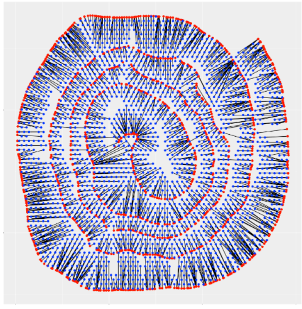

**b**

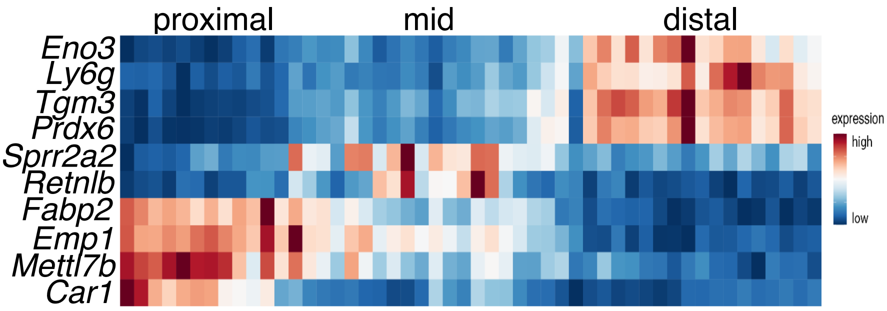

### Extended data Fig. 3

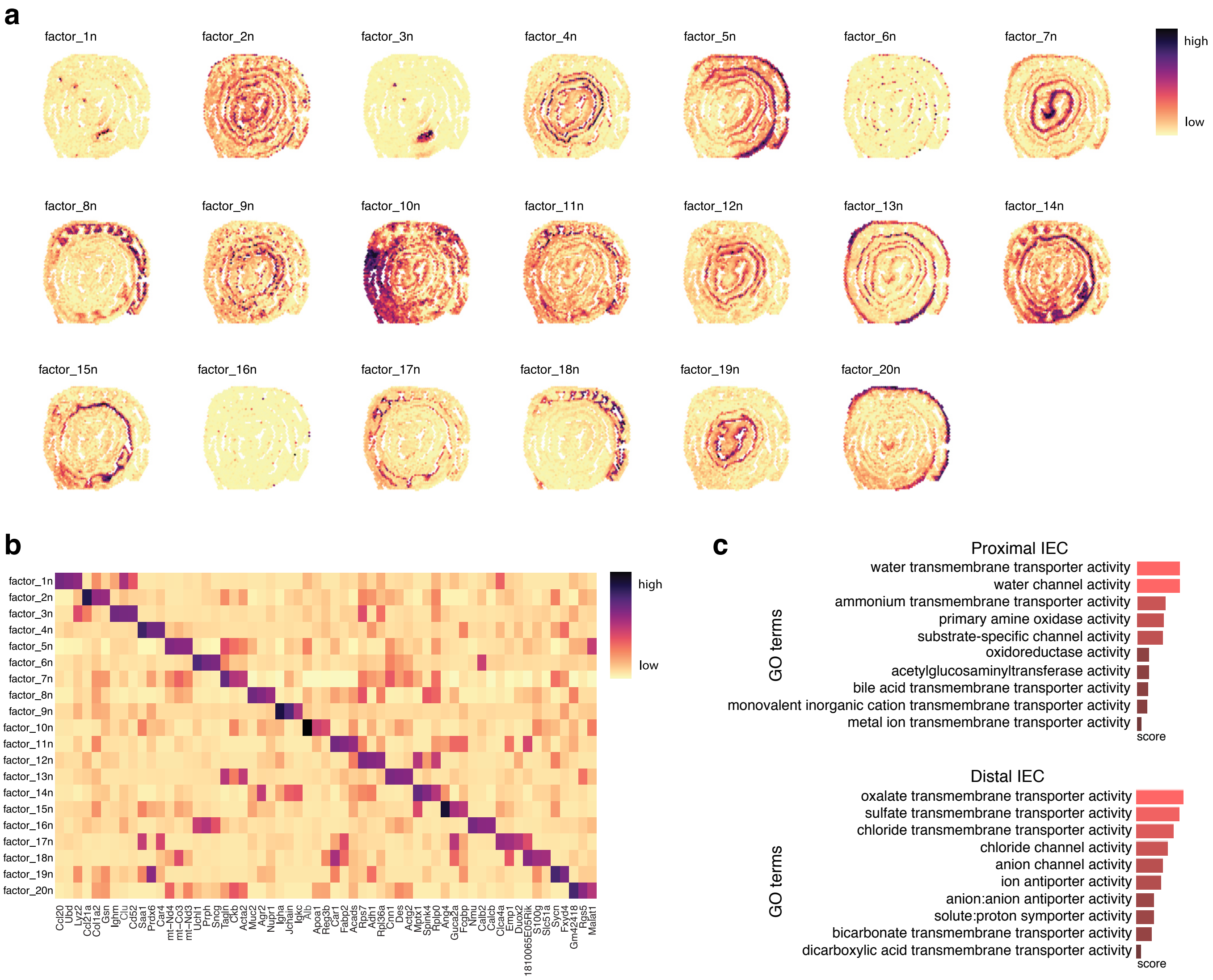

### Extended data Fig. 4

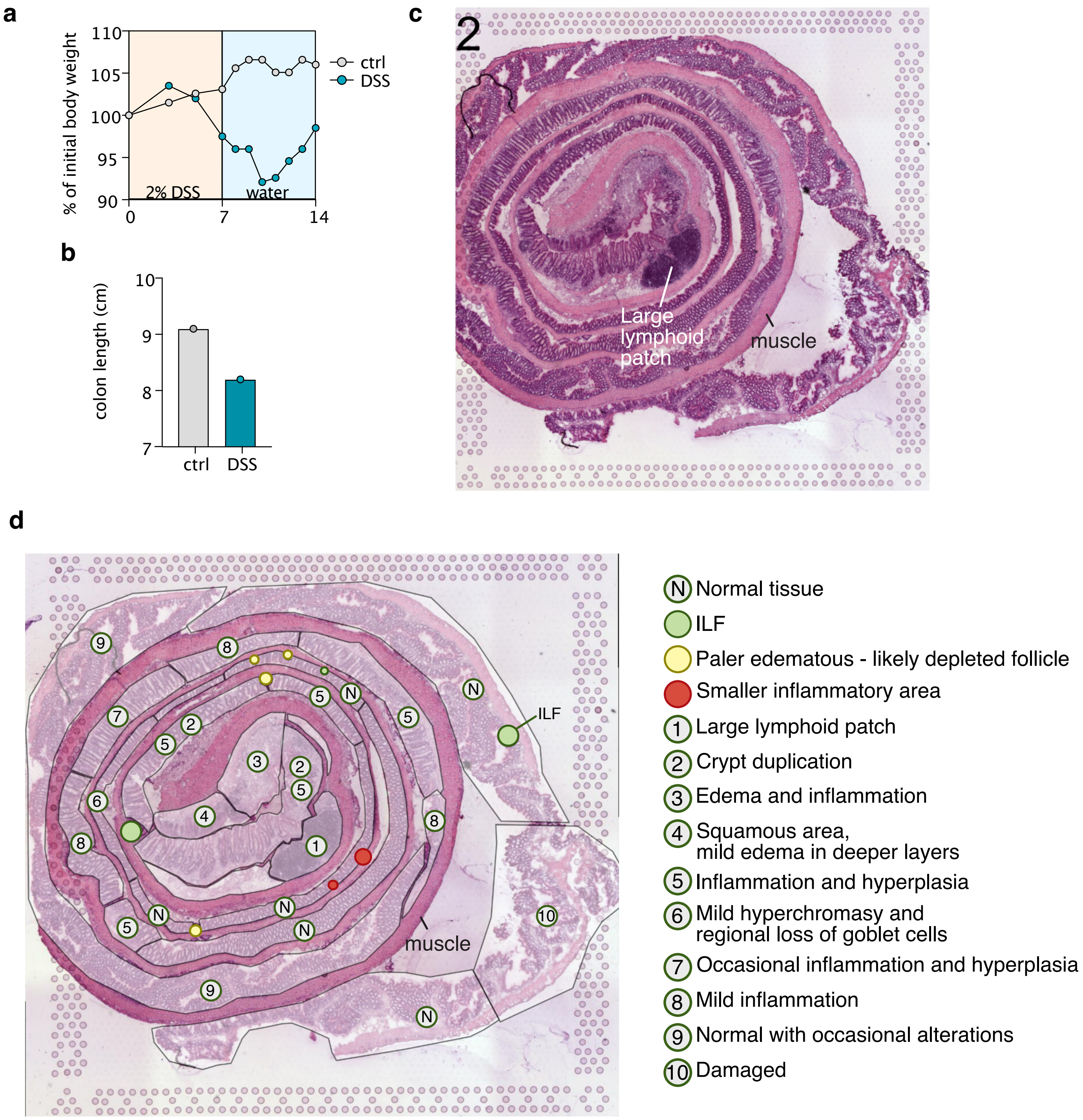

### Extended data Fig. 5

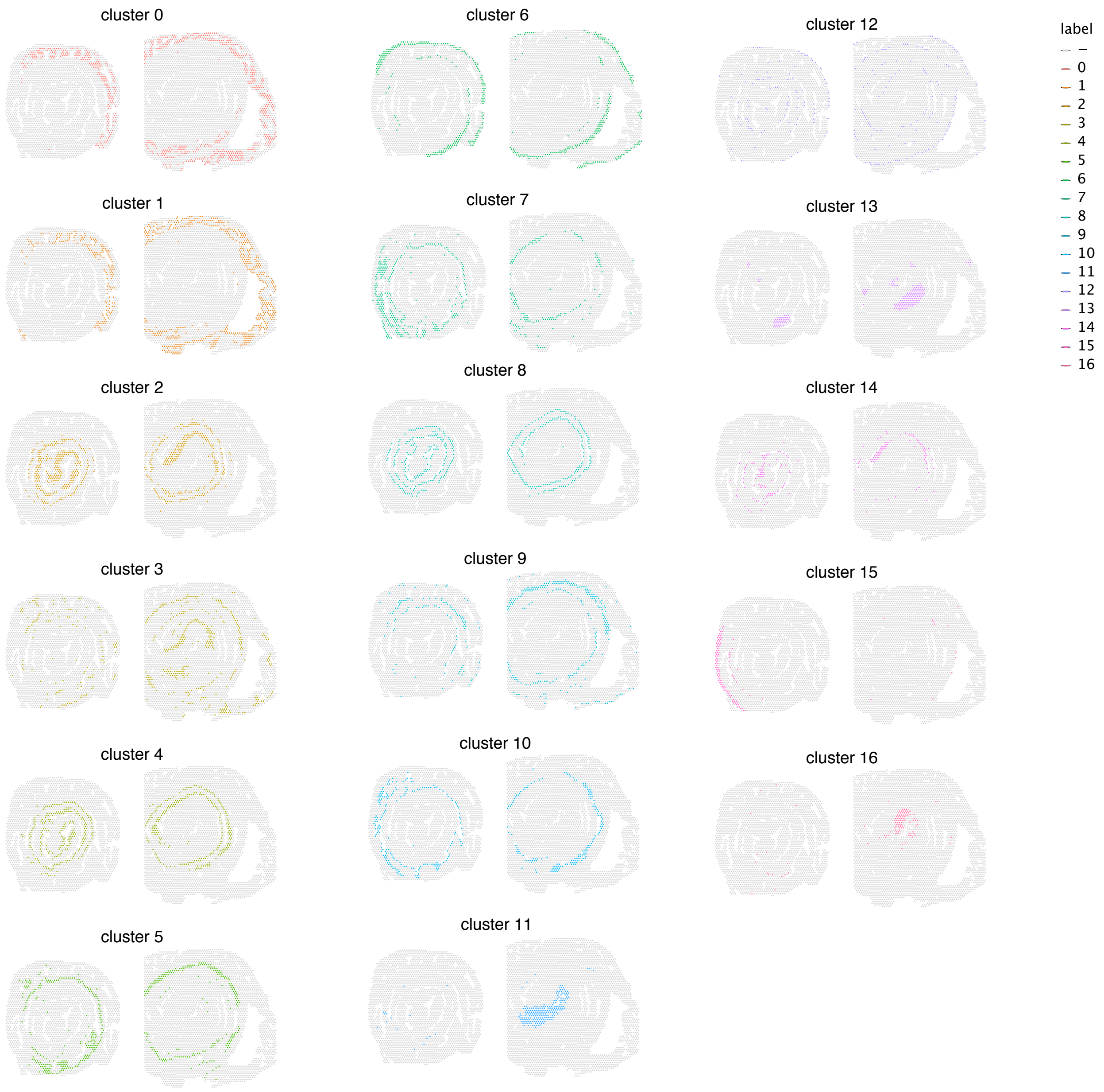

### Extended data Fig. 6

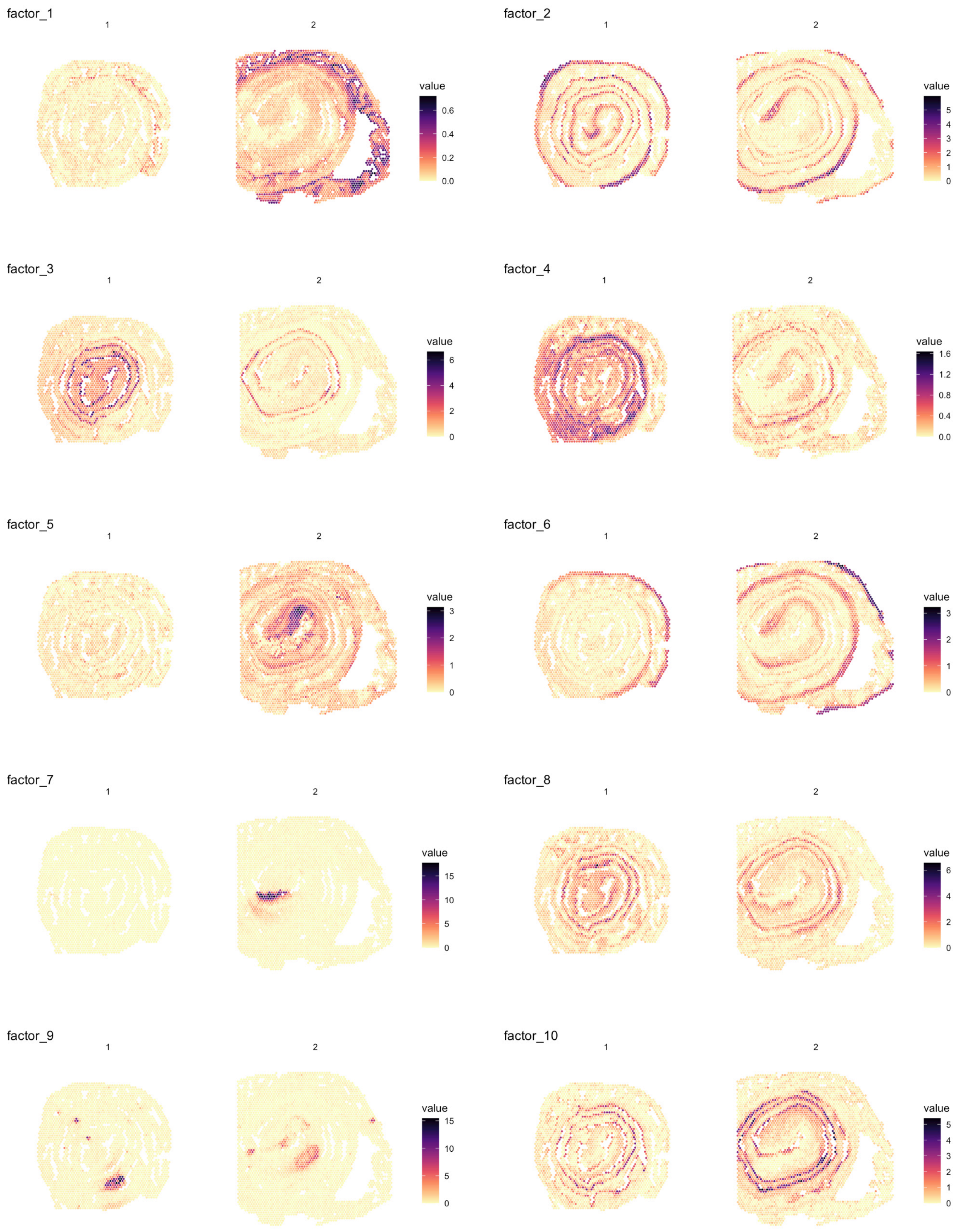

### Extended data Fig. 7

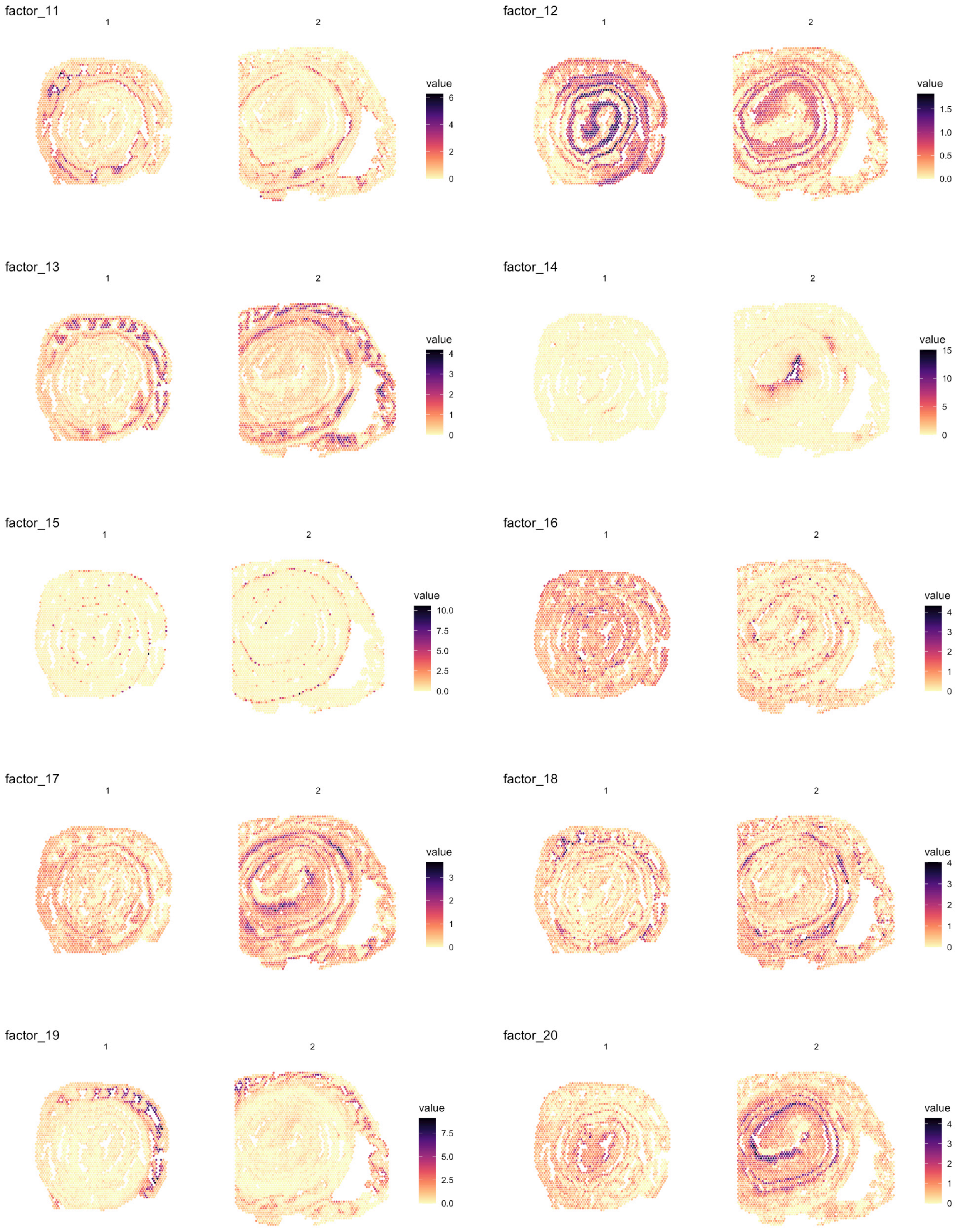

### Extended data Fig. 8

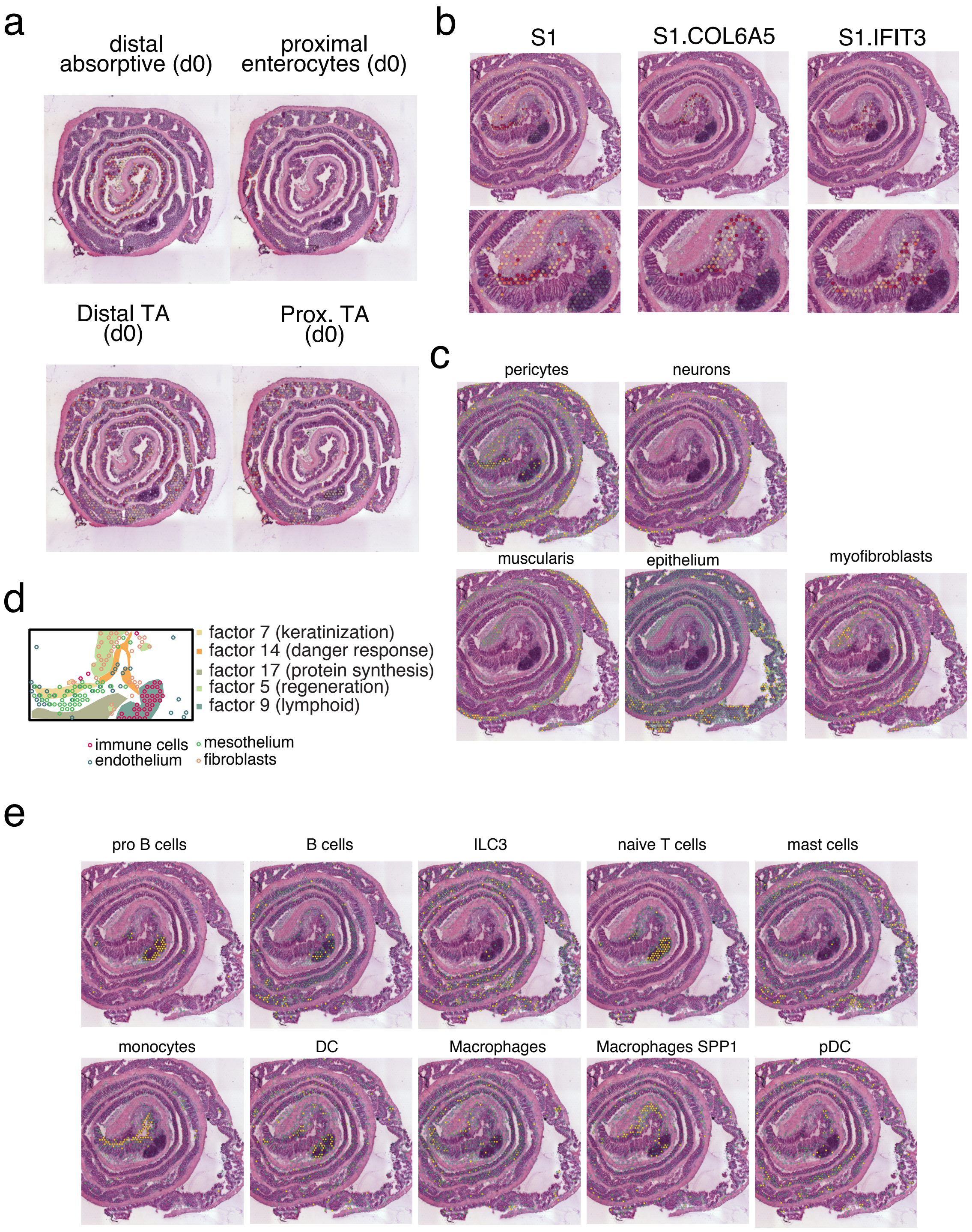

### Extended data Fig. 9

**a**

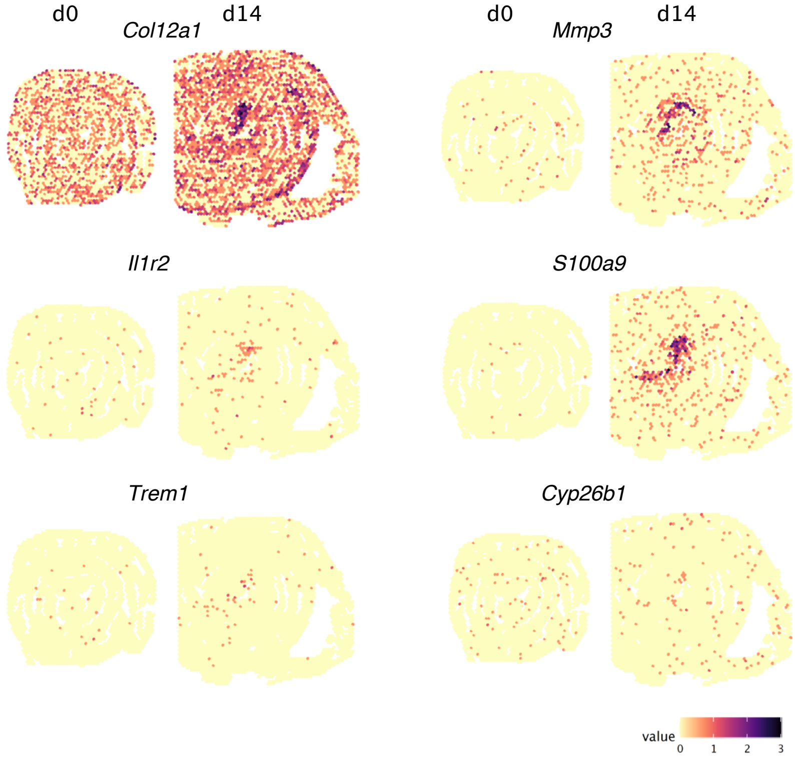


b

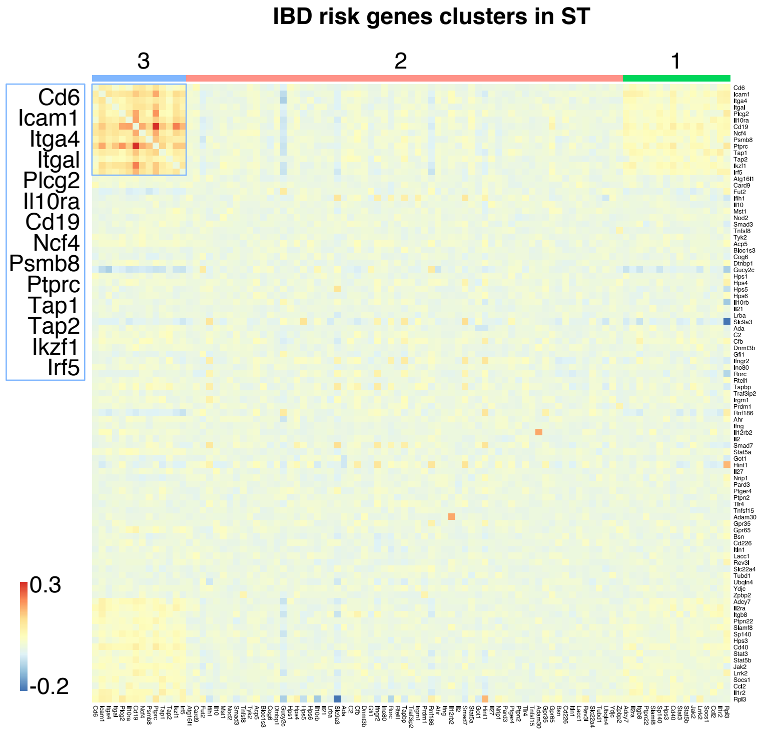
